# Disentangling signatures of selection before and after European colonization in Latin Americans

**DOI:** 10.1101/2021.11.15.467418

**Authors:** Javier Mendoza-Revilla, Juan Camilo Chacón-Duque, Macarena Fuentes-Guajardo, Louise Ormond, Ke Wang, Malena Hurtado, Valeria Villegas, Vanessa Granja, Victor Acuña-Alonzo, Claudia Jaramillo, William Arias, Rodrigo Barquera Lozano, Jorge Gómez-Valdés, Hugo Villamil-Ramírez, Caio C. Silva de Cerqueira, Keyla M. Badillo Rivera, Maria A. Nieves-Colón, Christopher R. Gignoux, Genevieve L. Wojcik, Andrés Moreno-Estrada, Tábita Hunemeier, Virginia Ramallo, Lavinia Schuler-Faccini, Rolando Gonzalez-José, Maria-Cátira Bortolini, Samuel Canizales-Quinteros, Carla Gallo, Giovanni Poletti, Gabriel Bedoya, Francisco Rothhammer, David Balding, Matteo Fumagalli, Kaustubh Adhikari, Andrés Ruiz-Linares, Garrett Hellenthal

## Abstract

Throughout human evolutionary history, large-scale migrations have led to intermixing (i.e., admixture) between previously separated human groups. While classical and recent work have shown that studying admixture can yield novel historical insights, the extent to which this process contributed to adaptation remains underexplored. Here, we introduce a novel statistical model, specific to admixed populations, that identifies loci under selection while determining whether the selection likely occurred post-admixture or prior to admixture in one of the ancestral source populations. Through extensive simulations we show that this method is able to detect selection, even in recently formed admixed populations, and to accurately differentiate between selection occurring in the ancestral or admixed population. We apply this method to genome-wide SNP data of ~4,000 individuals in five admixed Latin American cohorts from Brazil, Chile, Colombia, Mexico and Peru. Our approach replicates previous reports of selection in the HLA region that are consistent with selection post-admixture. We also report novel signals of selection in genomic regions spanning 47 genes, reinforcing many of these signals with an alternative, commonly-used local-ancestry-inference approach. These signals include several genes involved in immunity, which may reflect responses to endemic pathogens of the Americas and to the challenge of infectious disease brought by European contact. In addition, some of the strongest signals inferred to be under selection in the Native American ancestral groups of modern Latin Americans overlap with genes implicated in energy metabolism phenotypes, plausibly reflecting adaptations to novel dietary sources available in the Americas.

## Introduction

Admixed populations offer a unique opportunity to detect recent selection. In the human lineage, genomic studies have demonstrated the pervasiveness of admixture events in the history of the vast majority of human populations (Patterson et al. 2012; Hellenthal et al. 2014; Lazaridis et al. 2014). By inferring the ancestral origins of particular genetic loci in the genomes of recently admixed individuals, recent studies have provided evidence that such admixture has facilitated the spread of adaptative genetic mutations in humans. Notable examples include the transfer of a protective allele in the Duffy blood group gene likely providing resistance to *Plasmodium vivax* malaria in Malagasy and Cape Verdeans from sub-Saharan Africans (Hodgson et al. 2014; Pierron et al. 2018; Hamid et al. 2021), and the transmission of the lactase persistence allele in the Fula pastoralists from Western Eurasians (Vicente et al. 2019).

An ideal setting in which to test whether and how admixture contributed to genetic adaptation is Latin America. The genetic make-up of present day Latin Americans stems mainly from three ancestral populations: indigenous Native Americans, Europeans (mainly from the Iberian Peninsula), and Sub-Saharan Africans (Wang et al. 2007; Moreno-Estrada et al. 2013; Moreno-Estrada et al. 2014; Homburger et al. 2015; Chacon-Duque et al. 2018; Luisi et al. 2020) that were brought together starting ~500 years ago. The admixed genomes of Latin Americans are thus the result of an intermixing process between human populations that had been evolving independently for tens-of-thousands of years and that were suddenly brought together in a new environment. In this new environment, the ancestral genomes were quickly subjected to novel pressures that were largely unfamiliar from where they firstly evolved. Therefore, the genomes of Latin Americans potentially harbor signals of both older adaptations present in each of the ancestral populations, and more recent adaptations attributable to beneficial variants, e.g. introduced from a particular ancestral population, increasing rapidly in frequency post-admixture. Motivated by this, several studies have explored the genomes of admixed Latin Americans for signatures of selection, for example focusing on events occurring since the admixture event (Tang et al. 2007; Basu et al. 2008; Ettinger et al. 2009; Guan 2014; Rishishwar et al. 2015; Deng et al. 2016; Zhou et al. 2016; Norris et al. 2020; Vicuna et al. 2020). These studies have relied on an approach similar to that of admixture mapping, where the ancestry of a genomic region in each admixed individual is assigned to a particular ancestral population, a technique known as local-ancestry-inference (LAI). Loci with significantly more inferred ancestry inherited from one ancestral population are assumed to have evolved under some form of selection (Tang et al. 2007).

In addition, the genetic make-up of Latin Americans offers the opportunity to detect selection in their ancestral populations, as large cohorts of Latin Americans can be leveraged to reconstruct genetic variation patterns in each source population. This is of particular use for exploring selection in Native Americans, since Native groups are currently underrepresented in genomic studies (Sirugo et al. 2019) and as a consequence only a few studies have centered on detecting adaptive signals of indigenous groups from the Americas. Such studies have identified strong selective signals at different genes, particularly at those related to immunity, highlighting the selective pressures that Native Americans were subjected to after they entered the continent (Lindo et al. 2018; Reynolds et al. 2019; Avila-Arcos et al. 2020).

With some exceptions (Cheng et al. 2021), these studies either limited their analyses to Latin Americans with high Native American ancestry or used LAI to infer loci in individuals that derive from a Native American source. However, such approaches may result in a reduction of statistical power due to removal of individuals with non-Native ancestry, inaccurate local ancestry estimation and/or through removing segments challenging to assign.

Here we present a novel statistical model that identifies loci that have undergone selection before or after an admixture event (which we refer to as pre- or post-admixture selection, respectively). In contrast to previous methods, this approach is based on allele frequencies and does not require assignments of local ancestry along the genome. We illustrate the utility of our new method by performing a selection scan in five Latin American cohorts collected as part from the CANDELA Consortium (Ruiz-Linares et al. 2014). Our results suggest that several loci have been subjected to natural selection in admixed Latin American populations, and in their ancestral populations, replicating many of these signals using LAI. Many of the putative selected SNPs are strongly associated to relevant phenotypes, or act as expression quantitative loci (eQTL) in relevant tissues, providing further evidence of their functional effect. Overall, our analyses highlight the usefulness of our method to detect signals of selection in admixed populations or their ancestral populations, and reveal novel candidate genes implicated in the adaptive history of groups from the American continent.

## Results

### Overview of AdaptMix

In part following Balding and Nichols (1995), and analogous to previous approaches (Long 1991; Mathieson et al. 2015; Cheng et al. 2021), our model AdaptMix assumes that, under neutrality, the allele frequencies of an admixed target population can be described using a beta-binomial model, with expected allele frequency equal to a mixture of sampled allele frequencies from a set of groups that act as surrogates to the admixing sources (fig. 1). In our case the admixed target population is a Latin American cohort, defined below, and we use three surrogate groups to represent Native American, European, and African admixing source populations. The mixture values are inferred a priori, e.g. using ADMIXTURE (Alexander et al. 2009) (fig. 1a) or SOURCEFIND (Chacon-Duque et al. 2018), as the average amount of ancestry that each admixed target individual matches to a set of reference populations. (The reference populations used by these programs may be the same as the surrogate populations, but they need not be as illustrated below.) We find the variance parameter that maximises the likelihood of this beta-binomial model across all SNPs. This variance term aims to limit the number of false-positives attributable to genetic drift in the target population following admixture and/or the use of inaccurate surrogates for the ancestral populations. Then, at each SNP, we calculate the probability of observing allele counts equal to or more extreme than those observed in the target population, hence providing a *P*-value testing the null hypothesis that the SNP is neutral (see Methods).

**Fig. 1.**
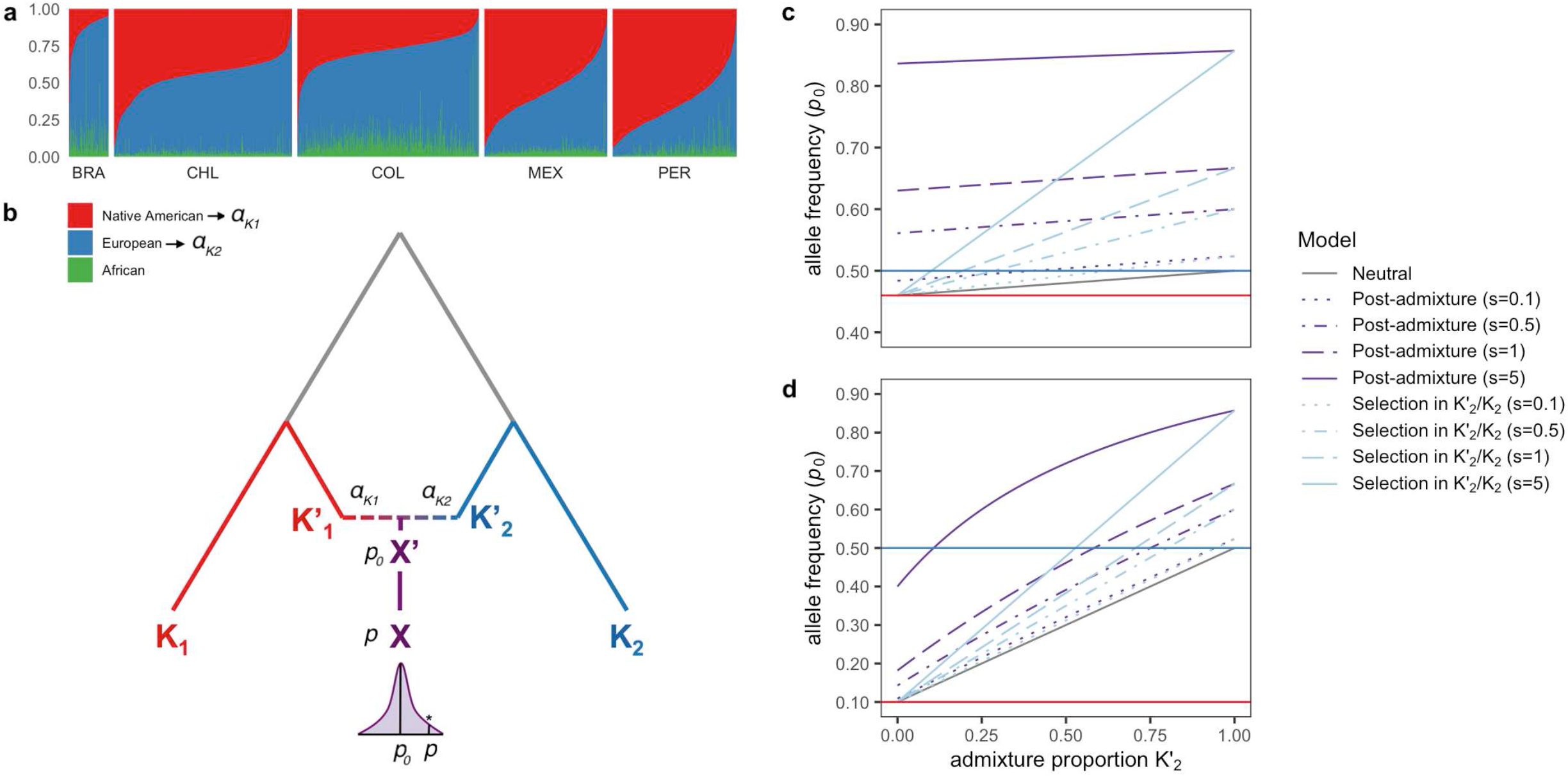
Schematic and intuition of the AdaptMix model. **(a)** For each CANDELA individual (columns), ADMIXTURE-inferred proportions of ancestry related to Native American, European, and African reference individuals. **(b)** Assuming only two admixing sources in this illustration for simplicity, the model assumes ancestral populations (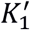 and 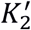) contribute ancestry proportions *α*_*K*_1__ and *α*_*K*_2__, respectively, to an admixed population (*X′*) that is ancestral to the tested population (A). Assuming neutrality, the expected allele frequency (*p*_0_) of *X′* is estimated using these proportions and the allele frequencies surrogate populations *K*_1_ and *K*_2_ related to 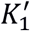 and 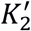, respectively. The sampled allele frequency (*p*) of *X* is compared to *p*_0_, with large deviations indicative of selection (shown with an asterisk in the distribution). **(c and d)** The relationship between *p*_0_, the expected allele frequency in the admixed population under neutrality or selection, and *α*_*K*_2__, the ancestry proportion contributed from ancestral population 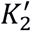. If selection occurred prior to admixture during the split between populations 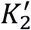 and its surrogate *K*_2_ (i.e. along the blue branch in **[a]**), this relationship increases linearly (blue lines), becoming more differentiated from neutrality (grey line) as the admixture from 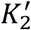 increases. In contrast, under selection post-admixture (i.e. along the purple branch in **a]**), the expected allele frequency (purple lines) can deviate from neutrality even when the admixture from 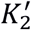 is near 0. The difference between the post-admixture and pre-admixture lines is more clear when allele frequencies in populations *K*_1_ and *K*_2_ are similar (top plot). Solid blue and red lines indicate the allele frequencies in the surrogate populations *K*_1_ and *K*_2_, which are used to calculate *p*_0_.

Assuming a pulse of admixture, this test is designed to detect selection occurring: (i) in the admixed population following the admixture event (e.g. along the purple line time period in fig. 1b), and/or (ii) in one (or more) of the source/surrogate pairings, i.e. following the split of the surrogate population from the admixing source it is representing (e.g. along the red and/or blue lines in fig. 1b). At SNPs with evidence of selection (i.e. low *P*-values), we distinguish between (i) and (ii) by exploring how genotype counts of admixed target individuals relate to their inferred admixture proportions contributed by each surrogate. Under scenario (i), we assume that selection affects all target individuals equally, regardless of their admixture proportions, which in turn assumes all ancestries were present when selection occurred. In contrast, under scenario (ii), we expect selection to more strongly affect one of the source/surrogate population pairings. Intuitively, if (ii) is true, individuals with nearly 100% ancestry from the source/surrogate pair experiencing selection will have genotype counts that deviate the most from expectations under the neutral model, while individuals with nearly 0% ancestry from this pair will have counts that closely follow the neutral model (fig. 1c). If instead (i) is true, this pattern is attenuated, though it can be challenging in practice to distinguish (ii) from (i) if allele frequencies strongly differ between surrogate groups (fig 1d). Assuming a multiplicative model of selection, we find the selection coefficients that maximize the fit of the data to model (i) and to model (ii) when separately treating each source/surrogate pair as the selected group. We report ratios of likelihoods, equivalent here to using differences in Akaike Information Criterion (AIC), to quantify our ability to distinguish among scenarios (i) and (ii). In summary, for each tested SNP we infer (a) a *P*-value testing the null hypothesis of neutrality, (b) the relative evidence (i.e. likelihood ratios) for whether selection occurred post-admixture or in one of the admixing sources and (c) the selection strength summed across time.

### Simulations

We tested our approach using simulations designed to resemble our Latin American cohort in terms of sample size, inferred admixture proportions, and the extent to which our surrogates match the true admixing sources (see Methods). At a false-positive rate of 5×10^-5^, these simulations indicate we have ~50-90% power to detect selection for scenario (i) (i.e., post-admixture selection) with selection strength (*s*) of 1.15-1.20 per generation in homozygotes carrying two copies of the selected allele, and selection occurring over 12 generations under various modes of selection (additive, dominant, multiplicative, recessive) (fig. 2a, supplementary fig. S1). For scenario (ii), in the case of selection occurring in the Native American source, power depends on the overall amount of Native American ancestry (fig. 2a). As an example, Brazil-like simulations (<15% average Native American ancestry) show little power, Colombia-like simulations (~30% average Native American ancestry) typically exhibit >50% power, and other simulated populations (~50–70% average Native American ancestry) exhibit >75% power under scenario (ii) assuming *s*=1.1 per generation over 50 generations, with similar power if instead s~1.025 over 150 generations (supplementary fig S2). Detecting selection occurring in the European source depends on the overall amount of European ancestry in a similar manner (e.g., fig. 2a, supplementary fig. S3). For SNPs where we detect selection, we mis-classify the type of selection ≤2% of the time, e.g., concluding post-admixture selection when the truth is selection in the Native American source ~1% of the time across all selection coefficients (fig. 2b). However, our approach often fails to classify selection scenarios unless selection strengths are large (e.g., *s*>1.1).

**Fig 2.**
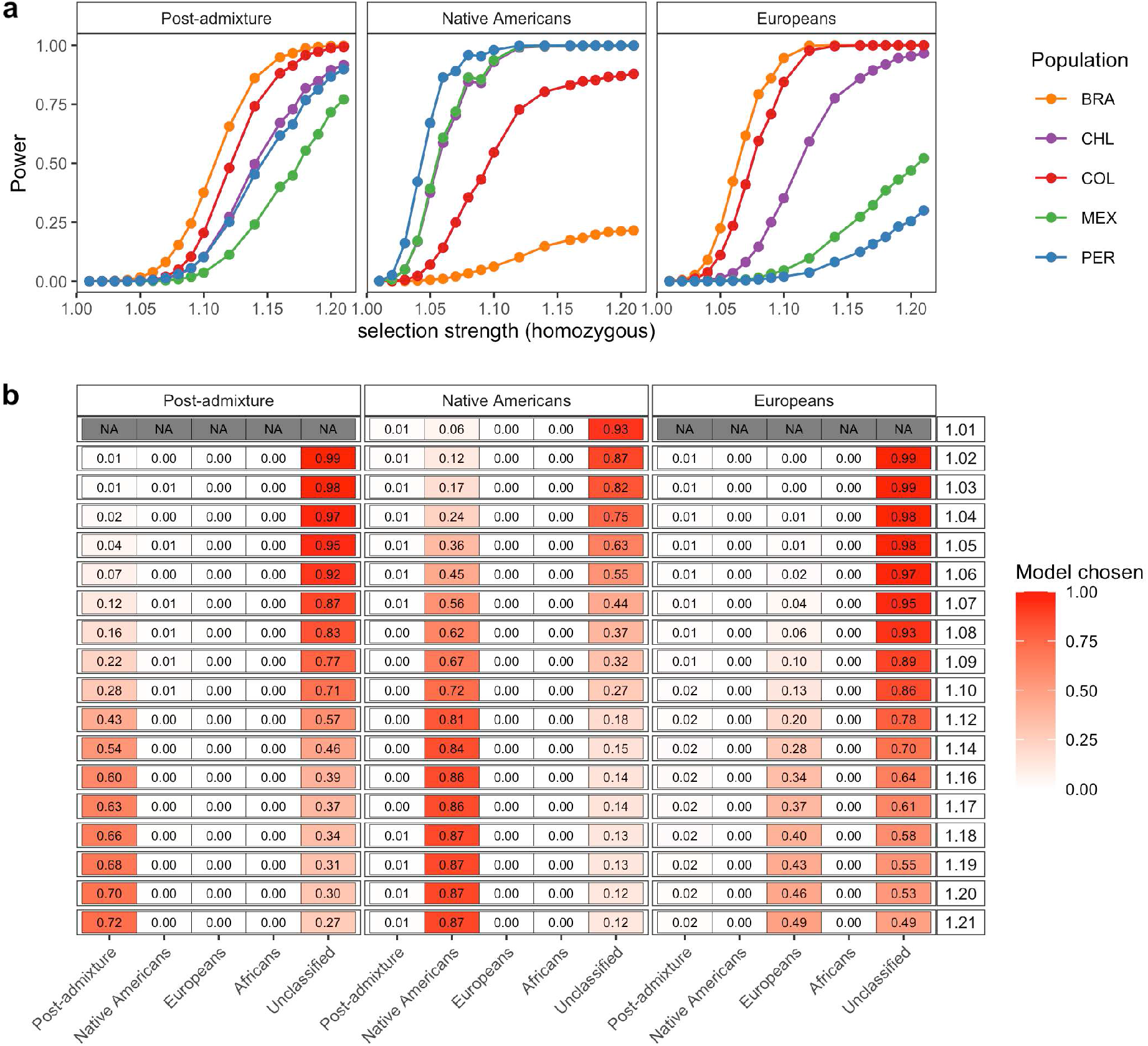
Performance of AdaptMix to detect and classify selection in simulated Latin American populations. **(a)** Power to detect selection post-admixture, selection in Native Americans, or selection in Europeans in simulated populations mimicking the Latin American cohorts. Power is based on a *P*-value cutoff that resulted in a false-positive rate of 5×10^-5^ in neutral simulations. The power estimated for a given selection coefficient is based on combining simulations using four different modes of selection (additive, dominant, multiplicative, recessive) over 12 generations for the post-admixture simulations, over 50 generations for the selection in Native American simulations, and over 25 generations for the selection in European simulations. Each simulation for a given combination of parameters consisted of 10,000 advantageous SNPs with a pre-selection minor allele frequency lower than 0.5. **(b)** The proportion of significant SNPs from (a) that were assigned to the correct simulated scenario of (left-to-right) post-admixture selection or selection in Native Americans or Europeans (using a likelihood ratio > 1,000 to make a call; otherwise ‘Unclassified’). Rows give the true selection coefficient (legend at right), and the heatmap values give the classification rate. Rows with N.A. shows instances with less than 50 selected SNPs for which the classification rate is not shown.

### Applying AdaptMix to the five Latin American cohorts of CANDELA

We divided Latin Americans into five cohorts based on country of origin: Brazil (n=190), Chile (n=896), Colombia (n=1125) Mexico (n=773), and Peru (n=834), using individuals sampled as part of the CANDELA Consortium (Ruiz-Linares et al. 2014), testing each cohort for selection separately (supplementary fig. S4). Analyzing each cohort by country of origin results in a higher number of individuals, and thus increases the statistical power to detect selection. As demonstrated in Chacon-Duque et al (2018), however, there is notable population sub-structure within each country. To test for robustness of our selection signals to this sub-structure, we supplemented each of these analyses by testing subsets of individuals within a country based on their inferred ancestry matching to Native American reference groups from Chacon-Duque et al. (2018). This gave six additional tested groups with sufficient ancestry represented: ‘Mapuche’ (n=434) in Chile, ‘Chibcha Paez’ (n=200) in Colombia, ‘Nahua’ (n=466) and ‘South Mexico’ (n=78) in Mexico, and ‘Andes Piedmont’ (n=195) and ‘Quechua’ (n=147) in Peru (supplementary fig. S5). To infer the proportion of African, European, and Native American ancestry in each Latin American, we applied unsupervised ADMIXTURE with *K*=3 clusters jointly to all CANDELA individuals and 553 Native American, Iberian, and West African reference individuals (fig. 1a).

Note that the choice of surrogate populations defines the selection test between each surrogate and its corresponding ancestral source in scenario (ii). In this way, our test acts as an analogue to *F_ST_* comparing two populations, but while accounting for admixture in one of the populations. As an illustration, we tested the Brazilian cohort for selection using northwest Europeans from England and Scotland (GBR) from the 1000 Genomes Project (1KGP) (The 1000 Genomes Project Consortium 2015) as a surrogate for the Brazilian cohort’s European ancestry source (supplementary fig. S6). Given the majority (~80%) of ancestry in the Brazilian cohort is related to Iberian Europeans, this test is most-powered to detect selection acting along the branch separating present-day northwest Europeans and descendants of Iberians who traveled to Brazil post-Columbus. In this analysis, we infer strongest signals of selection at the *HERC2/OCA2* and *LCT/MCM6* genes. This replicates previously reported selection signals when comparing northwest Europeans to present-day Iberians (Poulter et al. 2003; Bersaglieri et al. 2004), and likely indicates selection for lighter skin pigmentation and lactase persistence in northwest Europeans that is unrelated to any selection in the Americas. As another example, we also tested each Latin American cohort separately while using Han Chinese from Beijing (CHB) from the 1KGP as a surrogate for Native American ancestry (supplementary fig. S7). In this analysis, SNPs that follow model (ii) indicate selection along the branch separating present-day Han Chinese and Native American populations. For this test, we find the strongest signals of selection at previously reported selected genes in East Asians, including those related to alcohol metabolism such as *ADH7* and *ADH1B* (Galinsky et al. 2016; Gu et al. 2018) that both are classified as selection under model (ii). The strongest overall signal in this analysis overlapped the *POU2F3* gene, implicated in the regulation of viral transcription, keratinocyte differentiation and other cellular events, which has been reported to be under selection in Native American populations from throughout the Americas (Amorim et al. 2017).

For our main analyses, we use 205 Iberians (from 1KGP and Chacon-Duque et al. (2018)) to represent European ancestry surrogates. Therefore, given the likely short split time between present-day Iberians and Europeans that migrated to the Americas during the colonial era, we are underpowered to detect selection in the European source only (see simulations). We use 206 West Africans from the 1KGP to represent the African ancestry source, which has been reported as a good proxy to the African genetic sources (from Chacon-Duque et al. (2018)). For this reason, we should similarly have low power to find selection occurring only in the African source/surrogate. At any rate we do not test for selection related to African ancestry, because the Latin American cohort here have ~6% African ancestry on average, limiting power further. We combined 142 individuals with <1% non-Native American inferred ancestry from 19 Native American groups (supplementary table S1) to represent the Native American surrogate. By using individuals sampled from geographically spread Native American groups as the Native American ancestry surrogate, we aim to identify regional selection signals experienced by some Native American groups but not others. We also expect to have the highest power when testing for selection type (ii) in Native Americans, as there is likely to be the most time separating this ‘average’ Native American surrogate and the admixing source of each regional Latin American cohort. To avoid confounding our inference, we excluded individuals with >1% inferred ancestry matching to surrogates other than Native Americans, Iberian Europeans, and West Africans using SOURCEFIND (Chacon-Duque et al. 2018). Also, since the time since admixture among these groups is relatively short in the CANDELA cohort (likely <15 generations ago), detecting selection post-admixture can only identify relatively strong selection signals (see simulations).

### AdaptMix identifies 47 regions of putative selection

For each Latin American cohort, we considered SNPs under selection as those having *P*-values less than the 5×10^-5^ false-positive threshold in the population-matched neutral simulations, which corresponds to a model-based *P*-value of 6.75×10^-6^–1.07×10^-7^ (supplementary table S2). For Chile, Colombia, Mexico and Peru, we report loci that pass these criteria both in the analysis of all individuals from that country and in at least one of three alternative analyses for that country that are designed to test for robustness to latent population structure (supplementary fig. S8). The first of these alternative analyses consisted of identifying signals of selection using AdaptMix on each of the six Native American subsets defined above (e.g., in either the ‘Andes Piedmont’ or ‘Quechua’ subset when testing for selection in Peruvians) (supplementary table S3). The other two alternative analyses were based on LAI. In particular we used ELAI (Guan 2014) to assign each genomic region of an admixed individual to a Native American, European, or African ancestral source. For the second alternative analysis, designed to test for post-admixture selection, we assessed whether the proportion of ancestry inferred from one of these three sources in a local region deviated substantially from the genome-wide average (supplementary table S4). For the third alternative analysis, designed to test for selection in the Native American source, we instead used the Population Branch Statistic (PBS) (Yi et al. 2010) to test for selection in one of the six Native

American subset groups defined above, using allele frequencies computed from LAI-inferred Native American segments from the subset of individuals representing that Native American group (see Methods) (supplementary fig. S5 and supplementary table S5).

Overall, we find 51 candidate regions to have evidence of positive or purifying selection passing the criteria above, 47 of which target protein-coding genes (supplementary table S6 and fig. 3). Four of these 47 candidate gene regions contain at least one SNP exhibiting strong evidence (likelihood ratio >1,000) of selection affecting all admixed individuals regardless of ancestry proportions, which we assume reflects post-admixture selection. Furthermore, 18 of these 47 regions exhibit strong evidence of selection containing at least one SNP (likelihood ratio >1,000) in the Native American source only. The 25 remaining candidate gene regions are unclassified into either type of selection (likelihood ratio ≤1,000).

**Fig. 3.**
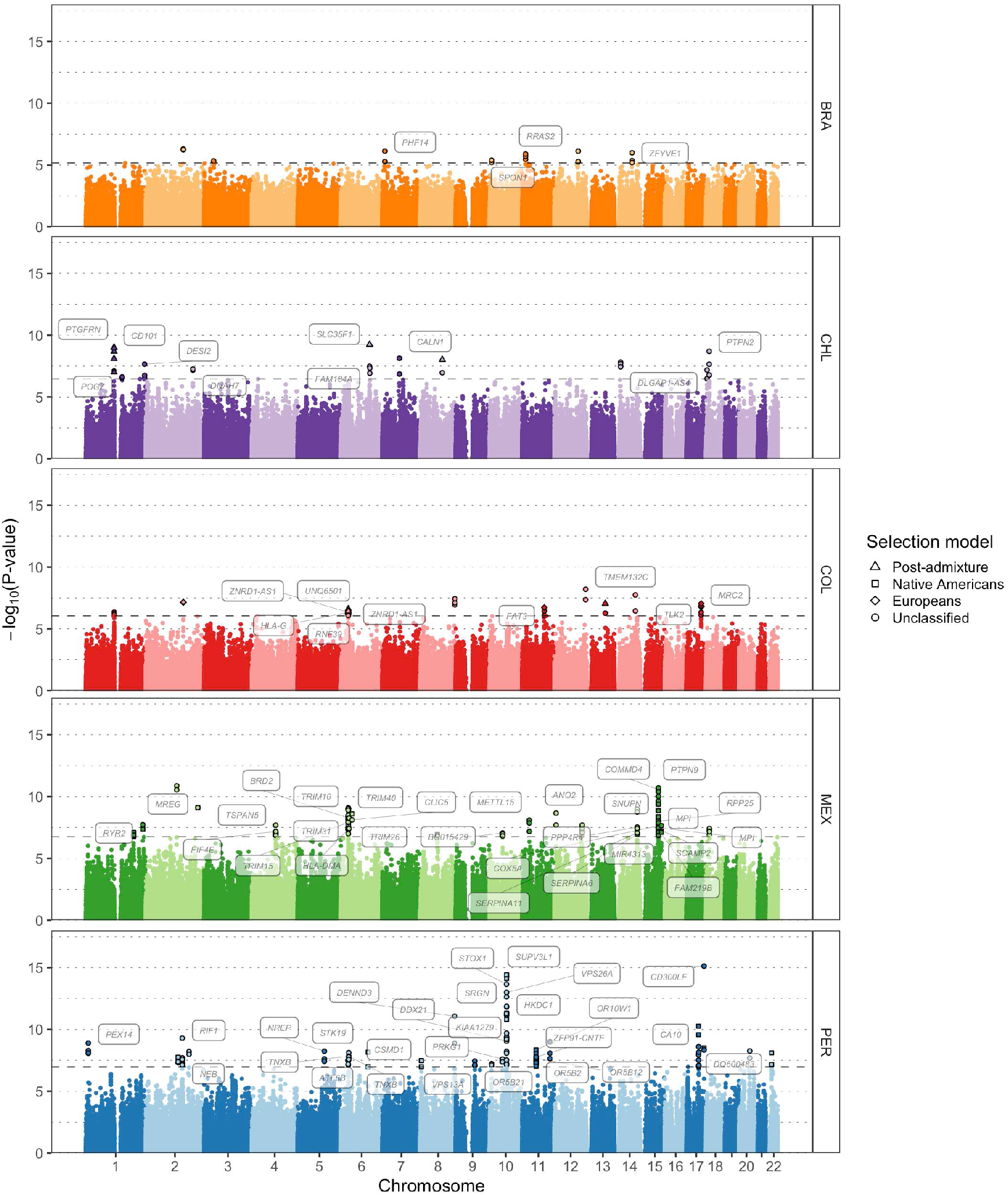
Genome-wide selection scan in five Latin American cohorts. Manhattan plot showing the genomic regions identified as selected via AdaptMix in each Latin American cohort. The dashed horizontal lines indicate the *P*-values cutoffs corresponding to a false-positive rate of 5×10^-5^ based on neutral simulations. Different shapes represent the most likely selection model. Names of genes associated with significant SNPs are shown.

To prioritize candidate casual genes, we annotated the protein-coding gene that had the highest overall Variant-to-Gene (V2G) scores (Ghoussaini et al. 2021) for the SNPs showing the strongest evidence of selection in each candidate gene region. The overall V2G score aggregates differentially weighted evidence of variant-gene association from several sources, including cis-QTL data, chromatin interaction experiments, *in silico* function predictions (e.g., Variant Effect Predictor from Ensembl), and distance between the variant and each gene’s canonical transcription starting site. For each of these candidate genes we then annotated the phenotype with the highest overall association score based on the Open Targets Platform (Koscielny et al. 2017).

While most of these associated phenotypes represent genetic disorders, syndromes, or different types of measurements (medically or non-medically-related), many are also related to immune response and diet – two major selective forces that shape the human genome (Karlsson et al. 2014; Fan et al. 2016). We therefore organize the description of our candidate selection signals into two main sections below that cover only these two features, with signals of all other hits in supplementary table S6. For brevity, below we only highlight putatively selected regions where at least one significant SNP had an associated GWAS or eQTL signal. For our significant SNPs related to immune-response genes, GWAS signals included SNPs associated to white blood cell counts in a large multi-continental cohort (including Latin American individuals) (Chen et al. 2020), and eQTL signals included cis-associated SNPs to gene expression in 15 immune-related cell types from the DICE project (Schmiedel et al. 2018). For our significant SNPs related to diet, GWAS signals included metabolic, anthropometric, and lipid levels from the UK Biobank cohort (Loh et al. 2018), and eQTL signals included cis-associated SNPs to gene expression in adipose, muscle, and liver tissue from the GTEx Project (Lonsdale et al. 2013).

### Signals at immune-related genes

Fifteen of the 47 candidate gene regions contained at least one protein-coding gene either related to the development or regulation of the immune system or that has been previously associated to the quantification of immune cell types, susceptibility progression to infectious diseases, or autoimmune disorders. For example, we replicate a well-known signal encompassing several immune-related genes at 6p21 that are part of the human leukocyte antigen (HLA) system (fig. 4 and supplementary fig. S9-S11). These included SNPs (AdaptMix *P*-value<5.00×10^-7^) near several MHC class I genes *(HLA-G, HLA-H, HLA-A*, and *HLA-J)* in each of the Chilean, Colombian, Mexican and Peruvian cohorts, with the Colombian cohort containing several SNPs classified as being selected post-admixture (likelihood ratio>1,000). Encouragingly, we inferred African ancestry enrichment (Z-score>2.5) in each cohort ~60kb downstream from our top AdaptMix signals using LAI, with maximum Z-score>9 (one-sided *P*-value<4.09×10^-21^) in the Chilean cohort (fig. 4). In addition, other signals were inferred upstream in the Chilean cohort at a 5’ UTR SNP of the *ZBTB12* gene (rs2844455, AdaptMix *P*-value=5.45×10^-8^), the Mexican cohort at an intronic SNP of *HLA-DMA* (rs28724903, AdaptMix *P*-value=3.87×10^-8^), and the Peruvian cohort at an intronic SNP of the MHC class III gene *STK19* (rs6941112, AdaptMix *P*-value=7.57×10^-9^). Many of these HLA genes have been previously characterized as subject to be under selection post-admixture in different Latin American populations by showing an excess of African ancestry at the HLA locus (Tang et al. 2007; Basu et al. 2008; Ettinger et al. 2009; Guan 2014; Rishishwar et al. 2015; Deng et al. 2016; Zhou et al. 2016; Norris et al. 2020; Vicuna et al. 2020).

**Fig 4.**
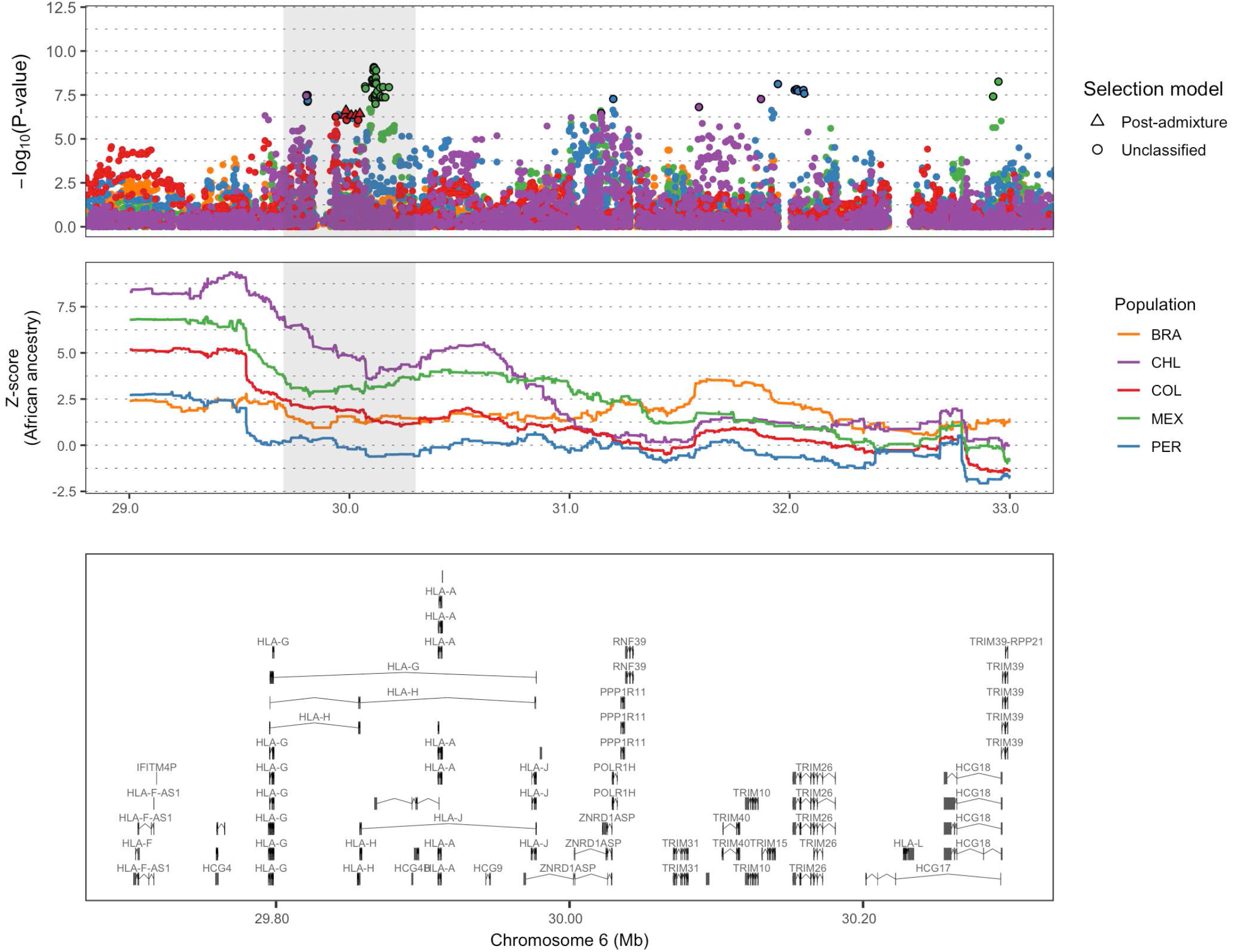
Regional selection plot at the HLA region in five Latin American cohorts. The top plot shows the –log_10_(*P*-values) of SNPs from AdaptMix, the middle plot shows *Z*-score values based on African local ancestry deviations, and the bottom plot shows genes in the region shaded in grey. Genomic coordinates are in Mb (build hg19 as reference) and genes shown include transcripts.

In addition to HLA, we infer previously unreported selection signals in four candidate gene regions that each harbor genes with well-established roles in the immune system, with each region containing at least one SNP significantly associated (*P*-value<5×10^-8^) to white blood cell counts or the expression of an immune-related gene in immune cells (*P*-value<10^-5^) (see Methods). Among these, one signal at 1p13 in the Chilean cohort encompasses the *CD101* gene (fig. 5a), which belongs to a family of cell-surface immunoglobulins superfamily proteins and plays a role as an inhibitor of T-cell proliferation (Soares et al. 1998; Bouloc et al. 2000). Within this region five SNPs are classified as being selected post-admixture and show also an increase of LAI-inferred European ancestry (maximum *Z*-score=3.40, one-sided *P*-value=3.36×10^-4^). Strikingly, the region contains a synonymous SNP (Ile588, CADD score of 9.23) (rs3736907, AdaptMix *P*-value=1.05×10^-9^) that strongly affects *CD101* expression in T cells (eQTL *P*-value < 2.42×10^-10^) and is associated with neutrophil (GWAS *P*-value=2.08×10^-10^) and total white cell count (GWAS *P*-value=3.61×10^-9^) (fig. 5a).

**Fig. 5.**
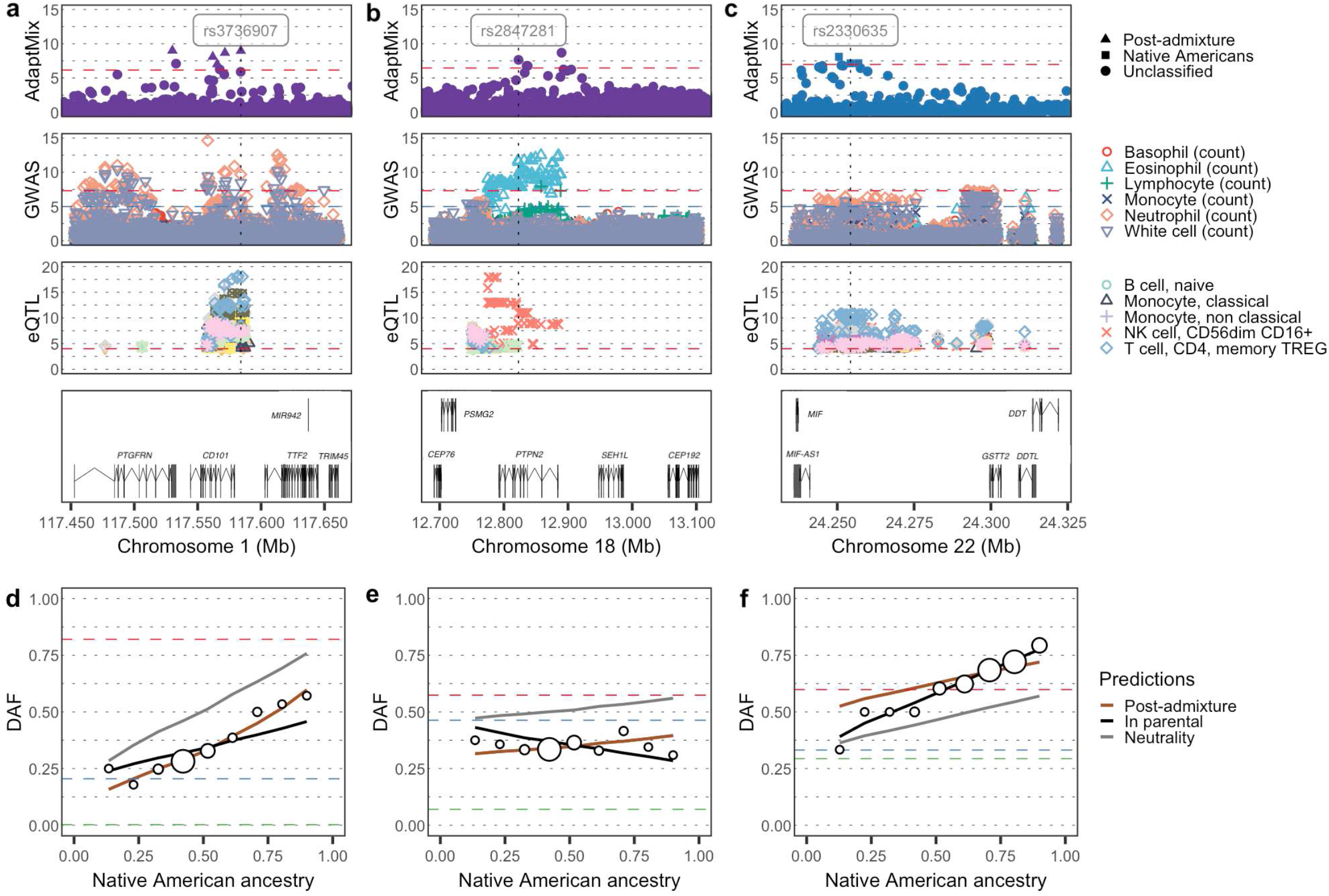
Genetic loci with signals of selection at immune-related genes. **(a)**, **(b)** and **(c)** Regional selection plot at three candidate regions of selection encompassing two immune-related genes in the Chilean and one immune-related gene in the Peruvian cohort, respectively. Each plot is composed of four panels (rows), consisting of –log_10_(*P*-values) of SNPs: (row 1) from AdaptMix; (row 2) associated with immune-related cell counts via GWAS (Chen et al 2020); (row 3) associated (as expression quantitative trait loci [eQTLs]) with expression of genes *CD101, PTPN2* and *MIF* for (a)-(c), respectively (Schmiedel et al. 2018); with (row 4) depicting genes in the region (in Mb, build hg19 as reference. Horizontal dashed lines give significance thresholds of (row 1) *P*-value =1×10^-5^ based on neutral simulations (row 2) *P*-value= 1×10^-5^ (blue line) and *P*-value = 5×10^-8^ (red line), and (row 3) *P*-value = 1×10^-4^. **(d)**, **(e)** and **(f)** Derived allele frequency (DAF) in admixed Latin Americans (white circles) stratified by proportion of inferred Native American ancestry, for the SNPs highlighted (vertical dashed line) in top row panels. The sizes of the circles are proportional to the number of individuals in that particular bin. Lines give expected DAF under neutrality (grey), post-admixture selection (brown) or selection in the Native source (black). Horizontal dashed red, blue, and green lines depict DAF for surrogates to Native American, European, and African sources, respectively.

The second signal, at 18p11 also in Chileans, encompasses the *PTPN2* gene, a tyrosine-specific phosphatase involved in the Janus kinase (JAK)-signal transducer and activator of transcription (STAT) signaling pathway (fig. 5b). The JAK-STAT pathway has an important role in the control of immune responses, and dysregulation of this pathway is associated with various immune disorders (Shuai and Liu 2003). Several SNPs with low AdaptMix *P*-values (*P*-value<1.69×10^-7^) in the 18p11 region are also associated with eosinophil counts (GWAS *P*-value<1.13×10^-10^) and the expression of *PTPN2* in natural killer (NK) cells (eQTL *P*-value<1.14×10^-9^) (fig. 5b).

The other two novel signals, both in the Peruvian cohort, are consistent with selection in Native Americans only (likelihood-ratio>1,000). The first, at 17q25, contains the *CD300LF* gene that encodes for a membrane glycoprotein that contains an immunoglobulin domain, and which plays an important role in the maintenance of immune homeostasis by promoting macrophage-mediated efferocytosis (Borrego 2013). Notably, a 3’UTR SNP (rs9913698, AdaptMix *P*-value=3.11×10^-9^) is strongly associated with monocyte count (GWAS *P*-value=1.00×10^-33^), total white cell count (GWAS *P*-value=5.96×10^-24^), lymphocyte count (GWAS *P*-value=2.50×10^-19^), and neutrophil count (GWAS *P*-value=1.30×10^-9^) (supplementary fig. S12). The second signal is at 22q11 adjacent to the *MIF* gene (fig. 5c), which is implicated in macrophage function in host defense through the suppression of anti-inflammatory effects of glucocorticoids (Calandra and Roger 2003). Variants within *MIF* have been recently associated to rheumatoid arthritis in southern Mexican patients (Santoscoy-Ascencio et al. 2020). The SNP rs2330635 (AdaptMix *P*-value=7.06×10^-8^) is strongly associated to the expression of *MIF* in T-cells (eQTL *P*-value<8.63×10^-5^) and NK cells (eQTL *P*-value=5.77×10^-9^) and is also marginally associated to neutrophil counts (GWAS *P*-value=2.46×10^-6^) (fig. 5c).

Overall, these findings suggest that some of the most robust signals of adaptation in the Americas can be ascribed to immune-related selective pressures. These plausibly resulted from both the introduction of novel pathogens after European colonization and the endemic pathogens encountered by the first Native Americans during the initial peopling of the continent.

### Signals at genes related to diet

Among the 47 candidate regions, nine regions contained at least one protein-coding gene potentially related to dietary practices through their association with metabolism-related phenotypes or anthropometric-related measurements (supplementary table S6). Among these, we infer three previously unreported signals where at least one of the selected SNPs was associated to metabolic- or anthropometric-related phenotypes, or to the expression of the candidate gene in adipose, muscle, or liver tissue (see Methods). One of these three hits (rs4636058, AdaptMix *P*-value=5.70×10^-10^), at 6p22 in the Chilean cohort, is classified as being selected post-admixture and shows an increase of LAI-inferred European ancestry (*Z*-score=3.78, one-sided *P*-value=7.82×10^-4^). It is located at 6q22 and encompasses the *SLC35F1* gene, whose function is not known, though several studies have associated this gene with different measurements of cardiac function (Hoffmann et al. 2017; Warren et al. 2017; Giri et al. 2019). Notably, SNP rs4636058 is marginally associated to cholesterol levels (UKBB GWAS *P*-value=3.8×10^-4^) and body fat percentage (UKBB GWAS *P*-value=4.29×10^-4^). Another of these three hits, at 1q31 in the Mexican cohort, is consistent with selection in Native Americans (likelihood-ratio>1,000) (fig. 6a). The 1q31 signal includes an intronic SNP (rs1171148, AdaptMix *P*-value=2.31×10^-8^) of *BRINP3*, a gene associated to body mass index in studies across different human groups (Pulit et al. 2019; Zhu et al. 2020). Within this region, various SNPs are associated to different metabolic-related phenotypes, including the SNP rs1171148 that is associated with hip circumference (UKBB GWAS *P*-value=4.96×10^-8^) and marginally associated with body mass index (UKBB GWAS *P*-value=5.51×10^-5^) (fig. 6a).

**Fig. 6.**
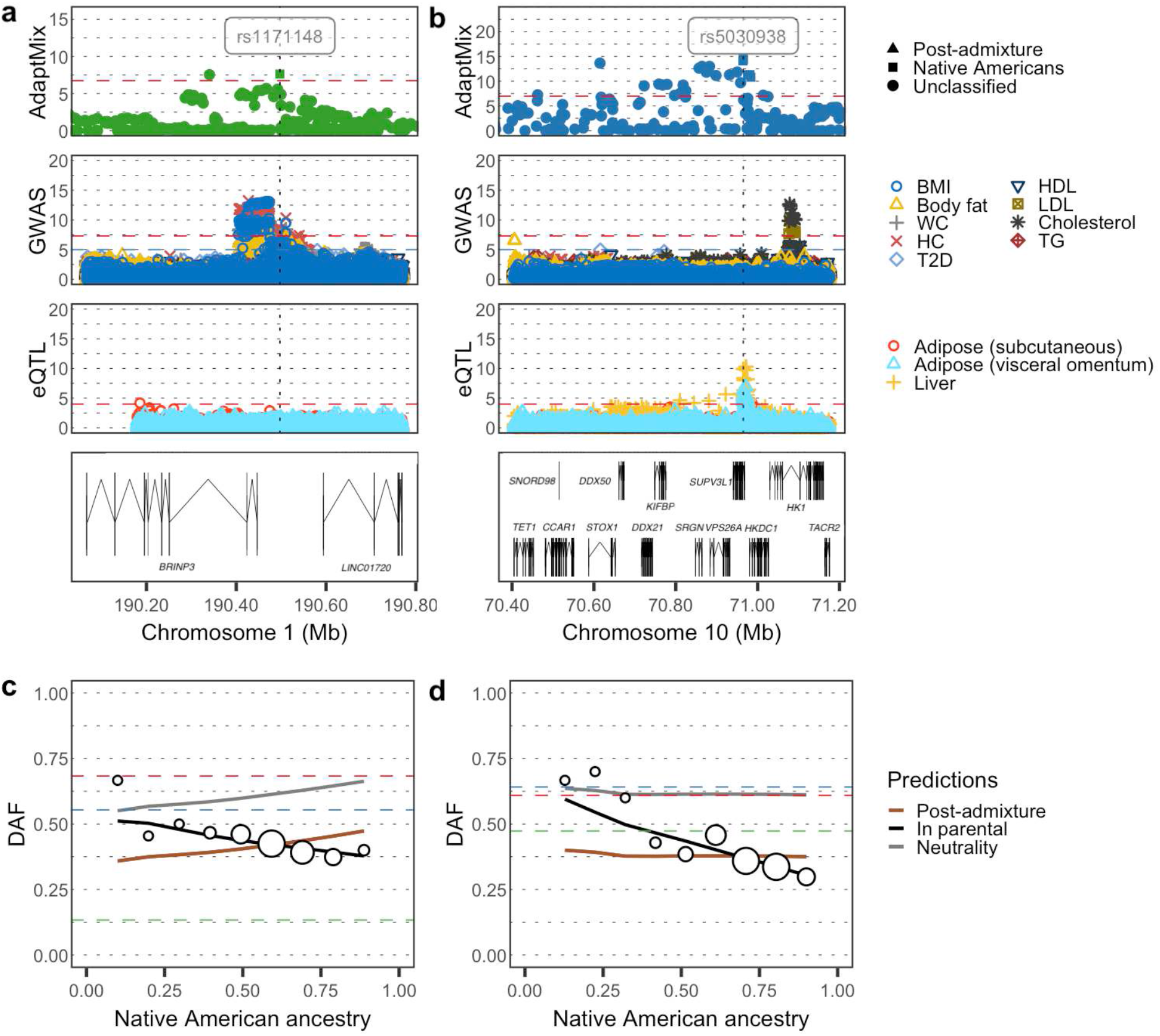
Genetic loci with signals of selection at metabolic-related genes. **(a)** and **(b)** Regional selection plot at two candidate regions of selection encompassing metabolic-related genes in the Mexican and Peruvian cohorts, respectively. Each plot is composed of four panels consisting of –log_10_(*P*-values) of SNPs: (row 1) from AdaptMix; (row 2) from the UK Biobank GWAS; (row 3) associated (as eQTLs) with expression of *BRINP3* and *HKDC1* for (a)-(b), respectively, (GTEx eQTL study); with (row 4) depicting genes in the region (in Mb, build hg19 as reference). Horizontal dashed lines give significance thresholds of (row 1) *P*-value= 1×10^-5^ based on neutral simulations (row 2) *P*-value = 1×10^-5^ (blue line) and *P*-value = 5×10^-8^ (red line), and (row 3) *P*-value= 1×10^-4^. **(c)** and **(d)** Derived allele frequency (DAF) in admixed Latin Americans (white circles) stratified by proportion of inferred Native American ancestry, for the SNPs highlighted (vertical dashed line) in top row panels. The sizes of the circles are proportional to the number of individuals in that particular bin. Lines give expected DAF under neutrality (grey), post-admixture selection (brown) or selection in the Native American source (black). Horizontal dashed red, blue, and green lines depict DAF for surrogates to Native American, European, and African sources, respectively.

Finally, the third hit (rs5030938, AdaptMix *P*-value= 3.79×10^-15^), which had the highest overall AdaptMix score, is inferred in the Peruvian cohort at 10q22 and indicates selection in Native Americans (likelihood-ratio>1,000) (fig. 6b). This SNP is associated with the expression of *HKDC1* in liver (eQTL *P*-value=2.19×10^-5^), adipose visceral (eQTL *P*-value=1.46×10^-5^), and adipose subcutaneous tissue (eQTL *P*-value=1.36×10^-4^) (fig. 6b). *HKDC1* encodes and hexokinase that catalyzes the rate-limiting and first obligatory step of glucose metabolism (Ludvik et al. 2016), and several studies have associated variants within this gene with glucose levels in pregnant women (Hayes et al. 2013; Guo et al. 2015; Kanthimathi et al. 2016; Tan et al. 2019) and with weight at birth (Warrington et al. 2019).

Overall, these results support previous hypothesis that genes related to energy metabolism were probably critical in the establishment of stable human populations in distinct ecoregions (Hancock et al. 2010), including those of the Americas (Amorim et al. 2017; Reynolds et al. 2019).

## Discussion

### Analytical considerations

Here we present AdaptMix, a novel statistical model that identifies loci under selection in admixed populations. Our model is based on the principle that allele frequencies in an admixed population can be modeled as a linear combination of the allele frequencies in the ancestral populations proportional to their admixing contributions, and that deviations from the expectation can be a product of selection. This selection test is related to the work of Long (1991) and Mathieson et al. (2015). One difference is that our approach directly infers and models the variance of the predicted allele frequencies in the admixed population given the set of surrogates used for ancestral sources. This parameter can help control for large deviations in allele frequency arising solely from genetic drift experienced in the admixed population (Long 1991; Bhatia et al. 2014) and/or from using inaccurate proxies for one or more of the source populations. In some applications here, e.g. the Brazilian cohort, AdaptMix gives *P*-values with a median near 0.5 as expected under the null hypothesis of neutrality, indicating a correction approach such as genomic control may not be necessary as in Mathieson et al. (2015) (supplementary fig. S13). However, simulations under neutrality that follow a slightly different model than our inference approach (see Methods), shows AdaptMix gives both an excess of high and low *P*-values relative to the uniform distribution expected under neutrality (supplementary fig. S14). This suggests our *P*-values are not well-calibrated, perhaps reflecting deviations from the underlying model and necessitating caution when choosing thresholds for significance. We thus based our significance thresholds on neutral simulations tailored to each cohort, and focus only on the strongest association signals that resulted in low false-positive rates based on simulated neutral SNPs. However, we caution that necessarily simulations are over-simplifications of complex latent demographic processes, and more work is required to verify these signals.

Another important contribution of our test is that it can infer whether selection disproportionately affects one source/surrogate pairing or affects all ancestry backgrounds equally. We assume selection affecting all ancestry backgrounds indicates selection occurring post-admixture, which is more parsimonious than an alternative explanation of independent selection events differentiating allele frequencies between each admixing source and its surrogate. For inferred selection in a source/surrogate pairing, this can reflect selection occurring in that source and/or its surrogate, possibly even following the admixture event. Post-admixture selection affecting only one source may be possible in cases of selection only occurring in a particular environment that is correlated with admixture fractions. For example, selection we detect to occur in Native Americans may be attributable to Europeans introducing a new environmental pressure (e.g. infectious disease) that disproportionately affected fitness in indigenous Americans. However, the split time between the true Native American ancestral source and our Native American surrogate is likely much longer than the time since colonial era admixture, suggesting selection pre-admixture as a more plausible explanation given the longer time to act. Supporting this, our inferred selection coefficients (which are summed over time) in cases where we conclude selection in Native Americans are typically greater than 2 (supplementary table S6). If selection had occurred post-admixture continuously over the last 12 generations (corresponding to an admixture date of ~1650CE), this value approximately corresponds to a per generation selection coefficient ~0.16, which is strong relative to previous reports of recent selection in human populations (e.g. Hamid et al. (2021)). In contrast, our four signals concluding post-admixture selection infer a per generation selection coefficient <0.1, which falls more in line with previous inference of selection strengths.

For 18 genomic regions where we conclude selection in the Native American source (supplementary table S6), it is possible this is capturing selection in (some subset of) groups that comprise the Native American surrogate group we use here, rather than in the (more localized) Native American source of the admixed population. The lack of overlap in selection signals when analysing the five CANDELA cohorts, and lack of concordance of our signals with those from PBS testing for selection in this combined Native American surrogate (supplementary fig. S15), suggests our signals are not being driven by selection in this combined population in practice. Furthermore, when using PBS to test for selection in LAI-inferred Native American segments from individuals with high degrees of ancestry recently related to the tested Native American source, an analysis that does not use the combined Native American surrogate, PBS scores for SNPs in 6 of these 18 regions fall into the top 99.99^th^ percentile (supplementary fig. S16-21), with the remaining 13 regions containing SNPs in the top 99^th^ percentile. However, relative to our approach, LAI-based inference (e.g., Avila-Arcos et al. (2020)) may be more robust to using combined data from multiple populations to represent one surrogate, since it only requires matching to a subset of individual’s haplotype patterns in the reference panel.

In general our approach has decreased power to distinguish whether selection occurred post-admixture versus in one of the ancestral sources, if reference population allele frequencies are very different and selection is weak (fig. 1c). Inferring excess ancestry matching using LAI would likely better capture post-admixture selection in such cases, e.g. a scenario where one population that is fixed (or nearly-fixed) for the protective allele intermixes with a population nearly-fixed for the non-protective allele, with the admixed population subsequently undergoing selection. An example of this is a recently reported excess of African ancestry, likely attributable to post-admixture selection, on the Duffy-null allele in inhabitants of Santiago Island in Cape Verde (Hamid et al. 2021). However, our test to detect whether *any* type of selection occurred should not be affected by these scenarios. In addition, our approach may identify post-admixture selection in scenarios that excess-ancestry LAI-based would miss by design, such as cases where the selected allele is at a similar frequency in all reference populations. Perhaps the most important contrast to LAI and other approaches detecting selection in admixed populations (Cheng et al. 2021), is that in principle our approach can be applied to populations that descend from the mixture of genetically similar groups, e.g. if using haplotype-based approaches (e.g. SOURCEFIND) to infer ancestry proportions. Future work should assess the power of this technique under such admixture settings.

While our method assumes a single pulse of admixture, theoretically our ability to diagnose and classify selection occurring in only one source should not be affected by multiple instances of (or continuous) admixture from that or any other source. This is because the signal of allele frequency deviation due to selection in such cases is entirely determined by the amount of ancestry inherited from that source, and not the number of admixture pulses. In contrast, if an admixed population experiences selection and then receives new migrants from one of the original admixing sources that are unaffected by this selection, e.g. later European migrants to the Americas, in theory this may attenuate our ability to determine that selection occurred post-admixture. However, in a simple scenario of one such additional admixture pulse, contributing 10-50% of DNA, the correct post-admixture selection theoretical model fits as well or better to the theoretical truth than does the incorrect model concluding selection in the source that did not contribute new migrants (supplementary fig. S22).

As noted above, and consistent with other tests comparing populations (Mathieson 2020), the choice of surrogate group can make a difference in the inferred selection signals. For example, our largest signal of Native American selection, at 10q22 and most strongly signalled in the ‘Andes Piedmont’ Peruvian subgroup, disappears if replacing the ‘combined Native American’ surrogate group with Han Chinese (CHB from the 1KGP) (supplementary fig. S7). In this case, the frequency of the putatively selected allele (rs5030938) is 67% in LAI-inferred Native haplotypes in the Peruvian ‘Andes Piedmont’ subgroup, which is notably higher than the 38-54% observed in LAI-inferred Native American haplotypes in four non-Peruvian sub-groups, and thus consistent with selection (supplementary table S7). However, it is lower than that of CHB (~76%,), which explains the lack of signal when using CHB as a surrogate. The frequency in Yakut, a Siberian group that perhaps better represents ancestral Native Americans than CHB does (Wang et al. 2007), is closer to that of frequency estimates across non-Peruvian Native American groups (0.46-0.5). In general, there is a trade-off between using surrogates more distantly related to the source, which may decrease power to find regional adaptation signals, versus choosing a more closely related surrogate, which may also decrease power by masking adaptation signatures that it shares with the target source (e.g. using Iberians as a surrogate for European ancestry of Latin Americans). Our inferred variance parameter can be used to investigate how well a given surrogate captures genetic variation in the target population, with for example the inferred variance using CHB as a surrogate ~5-10-fold higher relative to using the combined Native American surrogate.

### Selection signals detected in the CANDELA cohort

The candidate genes we infer to be affected by selection in Latin Americans and their Native American ancestors are best viewed in the context of other previously reported signals. Reynolds et al. (2019) recently performed a selection scan in three Native North American populations and identified some of the strongest signals at immune-related genes including the interleukin 1 receptor Type 1 (*IL1R1*) gene in a sample from several closely related communities in the southeastern United States, and the mucin 19 (*MUC19*) gene in a central Mexican population. We do not replicate the MUC19 signal in the CANDELA Mexican cohort, which could indicate that the Native American component in this cohort is not closely related to that of the central Mexican Native American group. Nonetheless, we found some of our strongest signals of selection at several loci encompassing genes involved in the immune response, including *CD300LF* and *MIF*, detected as being selected in the Native American ancestors of Peruvians. Interestingly, *CD300LF* promotes macrophage-mediated efferocytosis, while *MIF* play a role regulating macrophage function through the suppression of glucocorticoids. These observations suggest that these two genes might have perhaps evolved in a coordinated manner, possibly due to their phagocytic-related role against the novel pathogens encountered in the Americas.

Regarding signals of selection post-admixture, several studies have consistently shown adaptive signals in different Latin American populations at HLA by showing an excess of matching to African reference haplotypes using LAI (Tang et al. 2007; Basu et al. 2008; Ettinger et al. 2009; Guan 2014; Rishishwar et al. 2015; Deng et al. 2016; Zhou et al. 2016; Norris et al. 2020; Vicuna et al. 2020). Given that African ancestry was enriched at this region, the authors suggested that certain African alleles could have conferred a selective advantage to certain infectious diseases most likely brought by Europeans. While AdaptMix is only able to classify selection in one cohort (Colombia) out of our four HLA signals, we also replicated this excess of African ancestry in each of the CANDELA cohorts (supplementary fig. S9). There is some debate as to whether these signals are genuine or attributable to confounders such as inaccurate LAI inference (Pasaniuc et al. 2013). To illustrate the validity of these concerns, people with entirely Northwest European ancestry from Britain infer excess ancestry related to Africa in HLA, which – though perhaps influenced by genuine selection at HLA in Northwest Europeans – presumably does not reflect genuine recent African ancestry (supplementary fig. S23). Instead this is more likely attributable to the relatively high degree of genetic diversity in HLA mimicking African genetic diversity, illustrating how these LAI-based tests can give false-positive signals when testing for post-admixture selection. This may explain why AdaptMix does not replicate the moderate amount of excess African ancestry inferred by LAI at HLA in the Brazilian cohort (supplementary fig. S9), which is predominantly of European ancestry. Indeed regions under selection in admixed populations may be particularly difficult to classify accurately using LAI, e.g. with the HLA region here having the lowest overall LAI classification probability (supplementary fig. S24), especially in cases where the reference population have not experienced similar selection and hence may have poorly matching genetic variation patterns. As our approach does not require LAI, it is robust to these issues. While our model is not able to classify selection as post-admixture at most of our HLA signals, allele frequency patterns in the admixed cohorts are consistent with post-admixture selection and often show allele frequencies drifting away from those expected under our neutral model and towards those of the African or European reference population (supplementary fig. S25). This is most evident in the Colombian cohort, consistent with Africans contributing protective alleles as previously suggested (Tang et al. 2007; Basu et al. 2008; Ettinger et al. 2009; Guan 2014; Rishishwar et al. 2015; Deng et al. 2016; Zhou et al. 2016; Norris et al. 2020; Vicuna et al. 2020). In addition to HLA, we also identified a novel post-admixture selection signal in the Chilean cohort that was accompanied by a significant increase of European ancestry at the *CD101* locus, again, suggesting that protective alleles from Europeans might have also been adaptive to counter Old World-borne diseases brought to the Americas.

The signals encompassing genes related to metabolic and anthropometric-related phenotypes are consistent with novel dietary practices in the Americas driving adaptation, with many signals with an effect on relevant phenotypes and/or tissues, classified as being selected in the Native American source. Previous studies have shown evidence of adaptation at genes related to metabolic-related phenotypes and attributed the adaptation to dietary pressures in Native Americans. Avila-Arcos et al. (2020) recently reported strong signals of selection in the Mexican Huichol at several genes associated to lipid metabolism, including *APOA5* and *ABCG5*. We do not replicate these signals in the CANDELA Mexican cohort, which could indicate that the Native American component in this cohort is not closely related to that of the Huichol. The signals at *APOA5* and *ABCG5* are in line with a previous finding of a strong selection signal at another ATP-binding cassette transporter A1 (*ABCA1*) gene that has been associated with low high-density lipoprotein cholesterol in Latin Americans (Villarreal-Molina et al. 2008; Acuña-Alonzo et al. 2010). As the ABCA1 protein carrying the putative selected allele shows a decrease cholesterol efflux, the authors suggest that this variant could have favored intracellular cholesterol and energy storage, which in turn might have beneficially influenced the ability to accommodate fluctuations in energy supply during severe famines and during the regulation of reproductive function (Acuña-Alonzo et al. 2010). Lindo et al. (2018) used a genomic transect of Andean highlanders from northern Peru, and found the strongest signals of selection at *MGAM*, a gene related to starch digestion. The authors attributed this finding to a dietary-related selective pressure perhaps brought by the transition to agriculture in this region. AdaptMix shows evidence in the CANDELA Peruvian cohort within *MGAM* (rs7810984, AdaptMix *P*-value=1.79×10^-8^, above 99.9^th^ percentile) only when using CHB as a surrogate for Native American ancestry. This again illustrates how the choice of surrogate populations defines the selection test between each surrogate and its corresponding ancestral source. It is possible that by including Andean Native Americans in our Native American source population (supplementary table S1) we are affecting the power to detect selection in the Andean Native American ancestors of the CANDELA Peruvian cohort, analogous to how Lindo et al. (2018) no longer detect selection at *MGAM* if using PBS to compare ancient and present-day (Aymara) Andean groups.

Studies have also reported signals of selection in Native Americans groups shared with Siberian populations, which the authors interpreted as an adaptation to polyunsaturated-rich diets prior or close to the peopling of the Americas, likely in the Arctic Beringia. These included a signal overlapping the *WARS2* and *TBX15* genes, previously associated to body fat distribution and adipose tissue differentiation (Racimo et al. 2017), and the fatty acid desaturase (FADS) gene cluster that modulates the manufacture of polyunsaturated fatty acids (Amorim et al. 2017; Harris et al. 2019) (but see Mathieson (2020) for an alternative explanation of the *FADS* signal). Again, we inferred moderate selection evidence at these regions in the CANDELA Peruvian cohort only when using CHB as surrogate for Native American ancestry (SNP rs2361028 near *TBX15*, AdaptMix *P*-value=1.8×10^-7^, above 99.5^th^ percentile; SNP rs174576 within *FADS2*, AdaptMix *P*-value=3.8×10^-8^, above 99.5^th^ percentile). It is thus tempting to suggest that the three novel signals of selection AdaptMix classifies as being under selection in Native Americans might be related to dietary pressures experienced prior or during the peopling of the Americas (e.g., the *BRINP3* signal detected in Mexicans), or as a product for a greater reliance of domesticated crops including potatoes (3400–1,600 CE) (Rumold and Aldenderfer 2016) (e.g., the *HKDC1* signal detected in Peruvians). However, it is important to note that other factors may also be attributable for some of these selection signals.

Of potential adaptive interest is the *STOX1* gene detected in the Peruvian cohort close to our highest overall selection signal within *HKDC1* at 10q22 (fig. 6b). Mutations within *STOX1* have been associated to preeclampsia (Van Dijk et al. 2005; van Dijk and Oudejans 2011), a pathology of pregnancy characterized by high blood pressure and signs of damage to other organ system that can be lethal for the mother and for the fetus (Sibai 2003). Interestingly, in the single linkage study on preeclampsia conducted in Andean Peruvian families to date, SNPs within *STOX1* show marginal association (*P*-value=0.004678) (supplementary fig. S26) (Badillo Rivera and Nieves Colón et al. 2021). Given that high altitude is linked to an increased incidence of preeclampsia (Zamudio 2007), it is possible that natural selection has acted on genes related to this condition. Furthermore, the fact that variants within *HKDC1* are associated with glucose levels in pregnant women (Hayes et al. 2013; Guo et al. 2015; Kanthimathi et al. 2016; Tan et al. 2019) and considering the relationship between abnormal glucose levels and preeclampsia (Joffe et al. 1998; Weissgerber and Mudd 2015), it is also possible that natural selection has targeted variants at *HKDC1* due to its role in glucose metabolism.

Lastly, other environmental factors may also be attributable for some of these selection signals, such as infectious diseases. There is growing evidence of a link between metabolic diseases and innate immunity or inflammation (Pickup and Crook 1998; Kominsky et al. 2010; Lumeng and Saltiel 2011; Robbins et al. 2014). For instance, it has been shown that cholesterol plays a key role in various infectious processes such as the entry and replication of flaviviral infection (Osuna-Ramos et al. 2018). Additional studies in indigenous American populations will be needed to disentangle the putative selective pressures at these loci.

## Conclusion

We have presented a novel allele frequency-based method that identifies loci under selection in admixed populations, while determining whether the selection affected all ancestral sources equally, indicating selection following admixture, or in only one of the sources. The novel candidate genes under selection provide new insights into the adaptive traits necessary for the early habitation of the Americas and to respond to the challenge of infectious pathogens corresponding to European contact. Future functional investigations will allow a more detailed understanding of the consequences of selective pressures experienced in the American continent, including its effect on present-day health outcomes.

## Materials and Methods

### Genomic datasets

The Latin American individual samples analyzed here were part of CANDELA Consortium (Ruiz-Linares et al. 2014). The CANDELA Consortium samples (http://www.ucl.ac.uk/silva/candela) have been described in detail in previous publications (Ruiz Linares et al 2014; Chacon-Duque et al., 2018). These data include a total of 6,630 volunteers from five Latin American countries (Brazil, Chile, Colombia, Mexico and Peru). This dataset was genotyped on the Illumina HumanOmniExpress chip platform including 730,525 SNPs. We also collated reference populations from regions that have contributed to the admixture in Latin America. For Native American samples we used individuals previously genotyped by Chacon-Duque et al. (2018). This dataset compromises 19 Native American populations from throughout the Americas with genotype data (supplementary table S1). For all the analyses described, we have only retained Native American individuals that showed more than 99% Native American ancestry as estimated by ADMIXTURE (see below). For European samples, we used genotype data from Portuguese and Spanish, individuals previously genotyped by Chacon-Duque et al. (2018) and Spanish (IBS; Iberian Population in Spain) from the 1000 Genomes Project study (The 1000 Genomes Project Consortium 2015). For Sub-Saharan Africans, we used genotype data from Yoruba (YRI; Yoruba in Ibadan, Nigeria), and Luhya (LWK; Luhya in Webuye, Kenya) individuals from the 1KGP. The reference samples from Chacon-Duque et al. (2018) are described in more detail in the Supplementary Table 1 from the mentioned publication. For some of our analysis we also included the 103 Han Chinese from Beijing (CHB) and 85 Europeans from England and Scotland (GBR) from the 1KGP as a surrogate for the Native American and European source, respectively. Genotype data of the individuals from the 1KGP was downloaded from the 1000 Genomes Project FTP site available at ftp://ftp.1000genomes.ebi.ac.uk/vol1/ftp/.

### Data curation

We used PLINK v1.9 (Chang et al. 2015) to exclude SNPs and individuals with more than 5% missing data or that showed evidence of genetic relatedness as in Chacon-Duque et al. (2018). Due to the admixed nature of the Latin American samples, there is an inflation in Hardy-Weinberg *P*-values, and therefore we did not exclude SNPs based on Hardy-Weinberg deviation. After applying these filters, 625,787 autosomal SNPs and 7,986 individuals were retained for further analysis.

### Selecting admixed Latin American and reference individuals

In order to select admixed Latin American individuals (i.e. individuals with varying degrees of Native American, European and African ancestry), we conducted an unsupervised ADMIXTURE analysis at *K=3* using a set of 103,426 LD-pruned SNPs including Native Americans, Iberian Europeans and West Africans. We then removed non-admixed Latin American individuals that we define as having less than 10% or more than 90% Native American genome-wide ancestry. To avoid confounding our selection inference due to underlying population structure, we further excluded individuals with >1% inferred ancestry matching to surrogates other than Native Americans, Iberian Europeans, and West Africans using SOURCEFIND estimates obtained for the same individuals in Chacon-Duque et al. (2018). After this filtering procedure, the five Latin American populations consisted of 190 Brazilians (BRA), 1125 Colombians (COL), 896 Chileans (CHL), 773 Mexicans (MEX) and 834 Peruvians (PER). From our Native American, European, and Sub-Saharan African reference populations, we also removed individuals that contained more than 1% of ancestry from another group based on the ADMIXTURE analysis described above. After this extra filter our final reference dataset was composed of 142 Native Americans, 205 Europeans, and 206 Sub-Saharan Africans.

### Change in allele frequency under Wright-Fisher with multiplicative model of selection

Assuming a multiplicative model of selection and random mating, the frequency of the three genotypes in generation 1 at a biallelic locus with alleles A and a at frequencies *p* and 1 – *p*, respectively, in the previous generation is:

| AA | Aa | aa |
| --- | --- | --- |
| $(1 + s_1)^2 p^2 / c_1$ | $(1 + s_1) 2p(1 - p) / c_1$ | $(1 - p)^2 / c_1$ |

where *s*_1_ ∈ [−1, ∞] is the selection coefficient in generation 1 and *c*_1_ = (1 + *s*_1_)^2^*p*^2^ + (1 + *s*_1_)2*p*(1 – *p*) + (1 – *p*)^2^. Note that each copy of the A allele changes fitness by a factor of (1 + *s*_1_).

More generally, the allele frequency *p_g_* of allele *A* in generation *g* is:

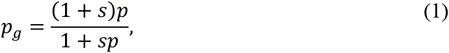

where

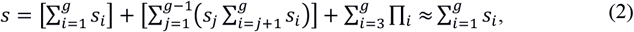

with *s_i_* the selection coefficient at generation *i* and *Π_i_* the sum of the products of all 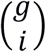 combinations of {*s*_1_, …, *s_i_*} values. The approximation in equation (4) assumes the s¿ are small, which should be a reasonable approximation based on e.g. estimated selection coefficients in humans.

### Testing for evidence of selection at a SNP

To assess the evidence of selection at a SNP, we employ a model inspired by that used in Mathieson et al. (2015) and based on the Balding-Nichols formulation (Balding and Nichols 1995). In particular for the allele count *X_j_* at SNP *j* in the target population, we assume:

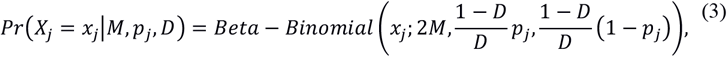

where *M* is the number of target individuals. The above model implicitly assumes that the frequency of the allele in the target population follows a *Beta*(*mean* = *p_j_, variance* = *D_p_j__*(1 – *p_j_*)). Under neutrality, we assume

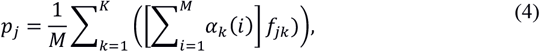

where *fjk* is the sampled frequency of the allele in the surrogate population at SNP *j* for source *k*, and ∝*_k_*(*i*) is the inferred admixture proportion from population *k* in individual *i*. We first find 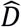 as the value of *D* that maximizes 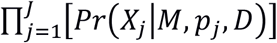, using the optim function in R with the ‘Nelder-Mead’ algorithm. Then, fixing 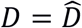 in equation (3), for each SNP we find the two-sided *P*-value testing the null hypothesis that the observed allele counts follow this neutral model.

The variance under (3) is small for SNPs with very high or very low *p_j_*, so such SNPs tend to reject this null model even in cases where the observed target population allele frequency does not deviate notably from its neutral expectation *p_j_* in (4). Therefore, we used an alternative parameterisation where we assumed the frequency of the allele in the target population follows a *Beta*(*mean* = *p_j_, variance* = *V*). This was achieved by substituting *D* in equation (3) at SNP *j* with 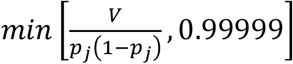, necessary to ensure numerical stability, and finding 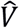. In practice this means that SNPs with minor allele frequency < (1.00001 × *V*) had variance (0.99999*p_j_*(1 – *p_j_*)) rather than *V*, though this approach gave sensible results in practice.

### Determining whether selection occurred pre or post-admixture

Consider the scenario in supplementary fig. S27, where sampled population C descends from an admixture of unsampled populations *A*\* and *B*\*, who are represented by sampled surrogate populations A and B, respectively. Our test aims to distinguish whether selection occurred postadmixture along branch (e) versus along any of branches (a)-(d). Let *f_c_* be the allele frequency of a sample from population C. At a neutral SNP:

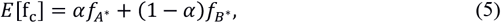

where *f_A*_* and *f_B*_* are true allele frequencies of *A*\* and *B*\* at the SNP, respectively, and *α* is the admixture proportion from *A*\*. Letting *f_k_* be the sampled allele frequency for population *k* serving as surrogate to the true admixing population *k*\*, it seems reasonable to assume:

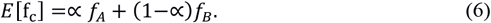

Note that this also holds under selection along branch (f) in supplementary fig. S27, which we ignore here (but which can be tested by comparing allele frequencies in *A* and *B*). Equation (6) assumes that *f_A_* and *f_B_* are equally good proxied for the admixing populations’ frequencies 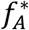 and *f_B*_*, respectively, at the SNP, which may not be true. We test the effect of this using simulations, described below, in which surrogates vary in how well they reflect their respective true admixing sources.

In the case of a multiplicative model of selection along branch (e) in supplementary fig. S27 at this SNP, using equation (1) we assume:

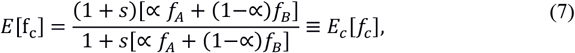

where s is the selection strength (i.e. equation [2]) along branch (e).

Alternatively, under a nultiplicative model for selection along branches (a) and/or (c) in supplementary fig. S27, with analogous results for selection along branches (d) and/or (b), instead we assume:

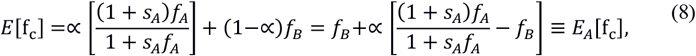

where *s_A_* is the selection strength along branches (a) and/or (c). Importantly, *E_A_*[*f_c_*] is linear in ∝, while *E_C_*[*f_c_*] is not, which we aim to exploit to distinguish between these two scenarios.

Here, assuming CANDELA population *T* can be described as a mixture of *K* sources, we assume the genotype *g_i_* of individual *i* ∈ [1, …, *M*] from *T* follows:

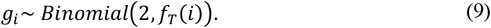

Under neutrality, we set *f_T_*(*i*) in (9) to:

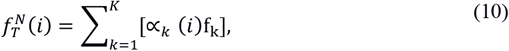

where *f_k_* is the sampled allele frequency at the given SNP for the surrogate population to the source contributing ∝*_k_*(*i*) admixture to individual *i*.

In the case of selection in *T* post-admixture, we generalize equation (7) and set *f_T_*(*i*) in (9) to:

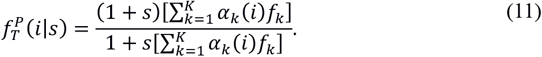

For the alternative case of selection along the branches separating source *A* and its sampled surrogate *A*\*, we generalize equation (8) and replace *f_T_*(*i*) in (9) with:

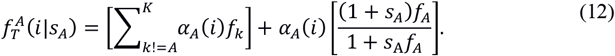

In practice, we fix ∝*_A_* (*i*) to be the proportion of DNA that each target individual *i* matches to surrogate *k* as inferred by ADMIXTURE. We define:

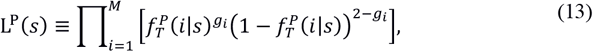

where *g_i_* is the genotype for target individual *i*. We use the optim function in R with the ‘Nelder-Mead’ algorithm to find the maximum-likelihood estimate (MLE) 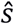, which is the value of s that maximizes equation (13).

Similarly we define:

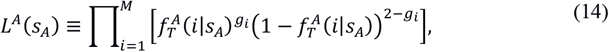

again finding 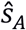, as the MLE for *s_A_*.

We note that 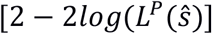 and 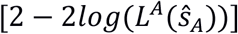 are analogous to AIC values for these respective models. Following AIC theory, we calculate:

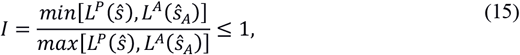

where, relative to the model with higher likelihood out of (13) and (14), the model with smaller likelihood is *I* times as probable to minimise the loss of information when used to represent the unknown true model (Akaike 1974).

Note we could analogously calculate the likelihood under the neutral model, i.e., using equation (10). Then, as an alternative to the selection testing approach described in Section ‘Testing for evidence of selection at a SNP’, we could use a likelihood-ratio-statistic approach to test for selection using either (13) or (14) as the alternative model likelihood. We explored this alternative testing approach, but do not use it here because it gave lower *P*-values when simulating under neutrality. This observation may in part be alleviated if we estimated *f_k*_* under both the neutral and alternative models rather than fixing *f_k*_* = *f_k_*. However, estimating *f_k*_* is confounded with estimating s or *s_A_* under the alternative models.

## Simulations

### Estimating how well each surrogate reflects its corresponding true admixing source

We aimed to generate simulations that mimic our real data. To do so, we first generate a measure of how well a sampled surrogate population *k* reflects its corresponding true (unknown) source population. In particular, we estimate a drift parameter *d_k_* in the following manner. First, at each SNP *j* we use nlminb in R to find the estimated values 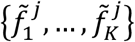 for {*f*_1*_, …, *f_K*_*}, respectively, that minimize:

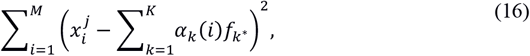

Where 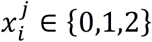 is the allele count for the admixed target individual *i* ∈ [1, …, *M*] at the SNP and each 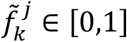. Then, for each source *k*, with observed allele counts 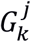 and total counts 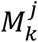 at SNP *j* in the surrogate population, following Balding-Nichols (Balding and Nichols 1995) we assume:

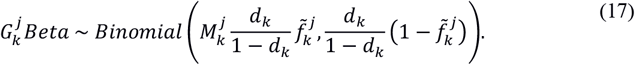

We then used the ‘Nelder-Mead’ algorithm in the optim function in R to find the *d_k_* ∈ [0, 1] that maximized the product of (17) across all SNPs. This gave the values reported in Table 1.

**Table 1.** Inferred *d_k_* measuring how well the sampled surrogate (column) reflect the true admixing sources for each target population (row).

| Target | Native American | European | African |
| --- | --- | --- | --- |
| Brazil | 0.173 | 0.007 | 0.102 |
| Chile | 0.02 | 0.011 | 0.226 |
| Colombia | 0.044 | 0.012 | 0.044 |
| Mexico | 0.024 | 0.007 | 0.223 |
| Peru | 0.015 | 0.009 | 0.119 |

Large estimated *d_k_* (>0.1) correspond to cases where there is little admixture from that source in our sampled individuals from that country, i.e. for African admixture in most countries and Native American admixture in Brazil. As values inferred using such little data are presumably unreliable, we cap them at 0.05 for the simulations below. While these values are a guide, in practice we adjusted these values by a multiple of 2-7 to generate neutral simulations that had the same inferred drift 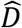, described in section ‘Testing for evidence of selection at a SNP’, as that observed in the real data.

### Generating simulated allele frequencies

We simulated admixed individuals who had experienced selection, with genome-wide admixture proportions ∝*_k_*(*i*) from source populations *k* ∈ [1, …, *K*] for simulated individuals *i* ∈ [1, …, *M*] matching those inferred by ADMIXTURE in the real data. To do so, for each simulation we repeated the following procedure:

1. For each source *k*, at each SNP we sample starting allele frequencies *f_k*_* from a 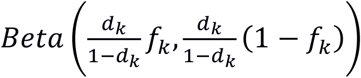, where *f_k_* is the sampled frequency of the respective surrogate population and *d_k_* are defined in Table 1 (but capped at 0.05).
2. We randomly select SNPs to undergo selection. If selection is occurring in source population fc prior to admixture, we randomly sample from among SNPs for which *f_k*_* < 0.5. If selection is occurring post-admixture, we instead randomly sample from among SNPs for which 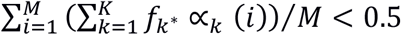.
3. We randomly select neutral SNPs from among all remaining SNPs, i.e., those not among the SNPs chosen in (2), in the real data.
4. To simulate selection:

- If selection is occurring prior to admixture, we simulate selection in the relevant source population for *g* generations under a specified model of selection (additive, dominant, multiplicative, recessive) using Wright-Fisher with a population size of *N_e_* indiviuals.
- If selection is occurring after admixture, we simulate selection separately in each of the source populations for *g* generations, under a specified model of selection using Wright-Fisher with a population size of *N_e_* individuals per population.
5. At each SNP, we sample allele counts for each individual *i* from a *Binomial*(2, *p_i_*) with 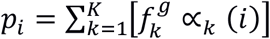 where:

- 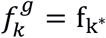 for neutral SNPs
- 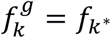 at selected SNPs for source populations *k* not undergoing selection (i.e., in cases where selection is pre-admixture)
- 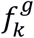is the sampled final frequencies in step (4) after *g* generations, at selected SNPs for source population *k* undergoing selection

We then analyse data from the simulated target population individuals using the real sampled data from the surrogate populations. For simulations here, we use *N_e_* = 10000 for the African, European, and Native American source groups.

Our procedure in steps (4)-(5) to simulate selection and admixture ensures the admixed individuals have variable admixture proportions while remaining computationally tractable. An alternative to this would be to generate *M* admixed populations using observed *f_k_* values, with the admixture proportions for population *i* equal to *α*_1_(*i*), …, *α_K_*(*i*), and then simulate each admixed population for *g* generations using Wright-Fisher, either with or without selection. Such simulations would match the approach used by our model to classify selection as type (i) or type (ii) (Section ‘Determining whether selection occurred pre- or post-admixture’). However, we chose the above for reasons of computational efficiency, as we have many individuals (i.e., *M* > 1000). Note also that our selection test (Section ‘Determining whether selection occurred pre- or post-admixture’) is different from this simulation procedure, in that our test models the combined allele frequency across all admixed individuals, using the mean admixture contributions across target individuals to calculate the expected frequency. This may explain why our model exhibits an excess of SNPs with small *P*-values even when simulating no selection. This is despite using all SNPs to infer our model’s variance parameter, which is designed to make more SNPs fit the model (likely explaining the excess of high *P*-values we also see, e.g., in supplementary fig. S14). While including this variance parameter does somewhat control *P*-values by e.g., giving a median *P*-value near 0.5, as expected under neutrality, our no-selection simulations suggest caution in directly using our model’s *P*-values for assessing selection evidence. This suggests some degree of plausible simulations would be helpful to calibrate the model’ s reported *P*-values.

### Local ancestry analysis

Local ancestry assignment was conducted using the HMM approach implemented in ELAI (Guan 2014). The phased genotype data needed as input was obtained by using SHAPEIT2 (Delaneau et al. 2012) with default parameter settings. Genetic distances were obtained from the HapMap Phase II genetic map build GRCh37 (Gibbs et al. 2003). As reference continental panels, we used the same Native American, European, and African individuals as in our AdaptMix analysis. ELAI was run setting the admixture generation parameter to 20, and with 20 rounds of EM iterations. To obtain local ancestry assignment probabilities, we conducted 10 independent runs and averaged probabilities across all runs as recommended in the ELAI manual. To test for local ancestry deviations we estimated *Z*-scores for each ancestry across each locus, and obtained the corresponding one-sided *P*-values testing for a positive deviation.

### Population Branch Statistic (PBS) analysis

We first selected Latin American individuals carrying a specific Native American ancestry component based on the inferred Native American ancestry proportions previously estimated by Chacon-Duque et al 2018 in the CANDELA sample. Specifically, for each Native American ancestry component, we selected CANDELA individuals with >10% inferred ancestry from that particular Native American ancestry component, and with <1% combined inferred ancestry combined across all other Native American components. Thus, each group of admixed Latin Americans was composed primarily of Native American ancestry from a particular Native American component, plus European and African ancestry. We then estimated allele frequencies for each Native American component by considering only alleles (i.e. haplotypes) that were considered of Native American origin with local-ancestry posterior probability >0.9. We only computed allele frequencies for a Native American component if all SNPs genome-wide had >100 alleles (haplotypes) assigned to Native American origin. This resulted in allele frequency estimates for six Native American components, including ‘Quechua’, ‘Andes Piedmont’, ‘Chibcha Paez’, ‘Nahua1’, ‘South Mexico’, and ‘Mapuche’ ancestral components (see Chacon-Duque et al. (2018) for a detail description of the inferred components). Pairwise *F_ST_* were then estimated using Hudson’s estimator as in equation 9 of Bhatia et al. (2013). The branch length (*T*) between two populations was computed as *T* = –log_10_(1 – *F_ST_*) (Cavalli-Sforza 1969). The Population Branch Statistic (PBS) (Yi et al. 2010) combines the pairwise branch lengths between three populations, which was computed as:

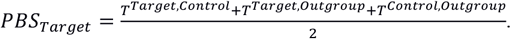

PBS values were computed for each Native American component, using all possible pairwise combinations of the other Native components as the control and outgroup populations. The rationale of this analysis was to try to find signals of selection exclusive to a given Native American group (i.e. that likely occurred after the divergence between Native American lineages). For some of our analysis we also used the CHB population from the 1000 Genomes Project, the European reference population, or the African reference population, as control and outgroup populations.

### Summary statistics for GWAS and eQTL data

To assess the biological consequence of selected variants, we queried summary statistics from GWASs of relevant phenotypes, and gene-expression data (i.e expression quantitative locus [eQTL] studies) from relevant cell or tissues. For our GWAS query, we retrieved data from immune and metabolic-related phenotypes, as these traits are known to have been subjected to strong selective pressures across several human groups (Fan et al. 2016). Immune-related phenotypes included (i) total white cell count, neutrophil count, lymphocyte count, monocyte count, basophil count, and eosinophil count from the Chen et al. (2020) GWAS study conducted across five continental ancestry groups. Metabolic-related phenotypes included body mass index (BMI), body fat percentage, type II diabetes status, hip circumference, waist circumference, HDL levels, LDL levels, cholesterol levels, and triglycerides levels (Loh et al. 2018). Summary statistics from these GWAS analyses were based on the UK BioBank cohort available at: http://www.nealelab.is/uk-biobank. For our eQTL query, we retrieved cis-associations summary statistics of 15 human immune cell types from the DICE (Database of Immune Cell Expression, Expression quantitative trait loci [eQTLs] and Epigenomics) project (Schmiedel et al. 2018), available at: https://dice-database.org/downloads. We also retrieved cis-association summary statistics from adipose (subcutaneous, and visceral omentum), muscle (skeletal), and liver tissue from the GTEx Project v7 (Lonsdale et al. 2013) available at: https://gtexportal.org/home/datasets.

## Acknowledgements

We thank the volunteers for their enthusiastic support for this research. We also thank Alvaro Alvarado, Mónica Ballesteros Romero, Ricardo Cebrecos, Miguel Ángel Contreras Sieck, Francisco de Ávila Becerril, Joyce De la Piedra, María Teresa Del Solar, Paola Everardo Martínez, William Flores, Martha Granados Riveros, Rosilene Paim, Ricardo Gunski, Sergeant João Felisberto Menezes Cavalheiro, Major Eugênio Correa de Souza Junior, Wendy Hart, Ilich Jafet Moreno, Paola León-Mimila, Francisco Quispealaya, Diana Rogel Diaz, Ruth Rojas, and Vanessa Sarabia, for assistance with volunteer recruitment, sample processing and data entry. We also thank Francois Balloux, Aida Andres, Mark McCarthy, and Etienne Patin for helpful discussion and critical comments on earlier versions of the manuscript. We are very grateful to the institutions that allowed the use of their facilities for the assessment of volunteers, including: Escuela Nacional de Antropología e Historia and Universidad Nacional Autónoma de México (México); Universidade Federal do Rio Grande do Sul (Brazil); 13° Companhia de Comunicações Mecanizada do Exército Brasileiro (Brazil); Pontificia Universidad Católica del Perú, Universidad de Lima and Universidad Nacional Mayor de San Marcos (Perú). Work leading to this publication received funded from: the National Natural Science Foundation of China (#31771393 to ARL), the Scientific and Technology Committee of Shanghai Municipality (18490750300 to ARL), Ministry of Science and Technology of China (2020YFE0201600 to ARL), Shanghai Municipal Science and Technology Major Project (2017SHZDZX01 to ARL) and the 111 Project (B13016 to ARL), the Leverhulme Trust (F/07 134/DF to ARL), BBSRC (BB/I021213/1 to ARL), the Excellence Initiative of Aix-Marseille University - A*MIDEX (a French “Investissements d’Avenir” programme to ARL), Wellcome Trust/Royal Society (098386/Z/12/Z to GH), the National Institute for Health Research University College London Hospitals Biomedical Research Centre, BBSRC (BB/R01356X/1), Universidad de Antioquia (CODI sostenibilidad de grupos 2013-2014 and MASO 2013-2014), Conselho Nacional de Desenvolvimento Científico e Tecnológico, Fundação de Amparo à Pesquisa do Estado do Rio Grande do Sul (Apoio a Núcleos de Excelência Program) and Fundação de Aperfeiçoamento de Pessoal de Nível Superior. JM-R was supported by a doctoral scholarship from CONCYTEC-PERU (224-2014-FONDECYT).

## Data availability

This project only analyses data that has been previously reported in other publications. Raw genotype data for reference populations can be accessed as described previously (The 1000 Genomes Project Consortium 2015; Chacon-Duque et al. 2018). Raw genotype data from CANDELA cannot be made available due to restrictions imposed by ethical approval. Summary statistics from the selection analysis will be deposited in a public repository upon publication.

## Software availability

Scripts for selection analyses will be uploaded to a software developer public repository upon publication. The current version of AdaptMix presented in this study is available upon request from.

## Notes

### Competing Interest Statement

The authors have declared no competing interest.

