## Supplementary online material for "Disentangling signatures of selection before and after European colonization in Latin Americans"

**Supplementary figures index**

**Supplementary figure S1** – Performance of AdaptMix to detect selection post-admixture in simulated Latin American populations
**Supplementary figure S2** – Performance of AdaptMix to detect selection in the Native American source population of simulated Latin American populations
**Supplementary figure S3** – Performance of AdaptMix to detect selection in the European source population of simulated Latin American populations
**Supplementary figure S4** – Map of sampled CANDELA individuals
**Supplementary figure S5** – Map of sampled CANDELA individuals based on their inferred ancestry matching to Native American reference groups
**Supplementary figure S6** – Manhattan plot in the Brazilian population using GBR from the 1000 Genomes Project as donor European population
**Supplementary figure S7** – Manhattan plots in the CANDELA cohort using CHB from the 1000 Genomes Project as donor Native American population
**Supplementary figure S8** – Flowchart of analyses
**Supplementary figure S9** – African local ancestry in Latin American populations
**Supplementary figure S10** – Native American local ancestry in Latin American populations
**Supplementary figure S11** – European local ancestry in Latin American populations
**Supplementary figure S12** – Regional selection plot at the *CD300LF* gene in the Peruvian population
**Supplementary figure S13** – Quantile-Quantile plot for AdaptMix in the CANDELA cohort
**Supplementary figure S14** – Distribution of *P*-values under neutrality in simulated Latin American populations
**Supplementary figure S15** – Correlation between PBS and AdaptMix scores in the CANDELA cohort
**Supplementary figure S16** – Regional AdaptMix selection plot at 1q31 in the Mexican population
**Supplementary figure S17** – Regional AdaptMix selection plot at 6p21 in the Mexican population
**Supplementary figure S18** – Regional AdaptMix selection plot at 15q24 in the Mexican population
**Supplementary figure S19** – Regional AdaptMix selection plot at 10q22 in the Peruvian population
**Supplementary figure S20** – Regional AdaptMix selection plot at 17q21 in the Peruvian population
**Supplementary figure S21** – Regional AdaptMix selection plot at 22q11 in the Peruvian population
**Supplementary figure S22** – Expected allele frequencies (green) at a SNP versus admixture received from source B, in a population undergoing selection following admixture between sources A and B
**Supplementary figure S23** – Local ancestry deviations in the GBR population from the 1000 Genomes Project
**Supplementary figure S24** – Mean posterior probabilities of local ancestry assignments in the CANDELA cohort and GBR population from the 1000 Genomes Project
**Supplementary figure S25** – Allele frequencies at significant SNPs at HLA loci

**Supplementary figure S26** – Selection signals in the Peruvian cohort and transmission disequilibrium test for preeclampsia at *STOX1*
**Supplementary figure S27** – An admixture graph showing the relationship between two parental populations, and an admixed population

**Supplementary tables index**

**Supplementary table S1 –** Reference population samples.
**Supplementary table S2** – Candidate SNPs under selection in the CANDELA cohort based on AdaptMix.
**Supplementary table S3** – Candidate SNPs under selection in CANDELA cohorts with Native American ancestry from a specic Native American groups based on AdaptMix.
**Supplementary table S4** – Candidate SNPs under selection in the CANDELA cohort based on local ancestry deviations (LAI).
**Supplementary table S5** – Candidate SNPs under selection based on PBS analysis in CANDELA cohorts with Native American ancestry from a specific Native group.
**Supplementary table S6** –Candidate SNPs under selection in the CANDELA cohort robust to popualation structure based on AdaptMix.
**Supplementary table S7** – Candidate SNPs under selection in the CANDELA cohort robust to popualation structure based on AdaptMix with associated derived allele frequencies.

Supplementary tables S1 to S7 as a separate Excel file.

### Supplementary figures

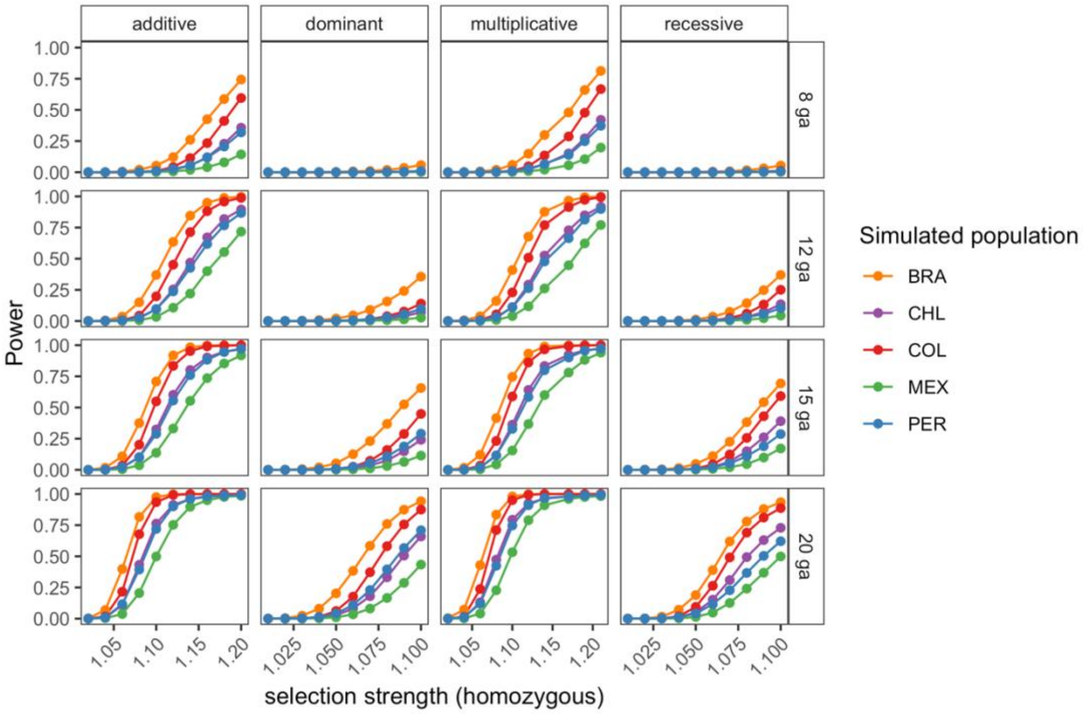


**Supplementary figure S1. Performance of AdaptMix to detect selection post-admixture in simulated Latin American populations.** Power to detect selection post-admixture in simulated admixed Latin American populations under three different admixture dates (rows), and four different selection models (columns). The power is based on a *P*-value cutoff that resulted in a $5 \times 10^{-5}$ false-positive rate in neutral simulations. Simulation parameters including sample sizes, inferred admixture proportions, estimated extent to which each ancestral source is captured by its corresponding surrogate population, and estimated mixture model fit are matched to those observed in the real data. Each simulation for a given combination of parameters consisted of 10,000 advantageous SNPs with minor allele frequency lower than 0.5. Note that the x-axis for the “dominant” and “recessive” models is half that of the “additive” and “multiplicative” models; power is broadly similar under each model.

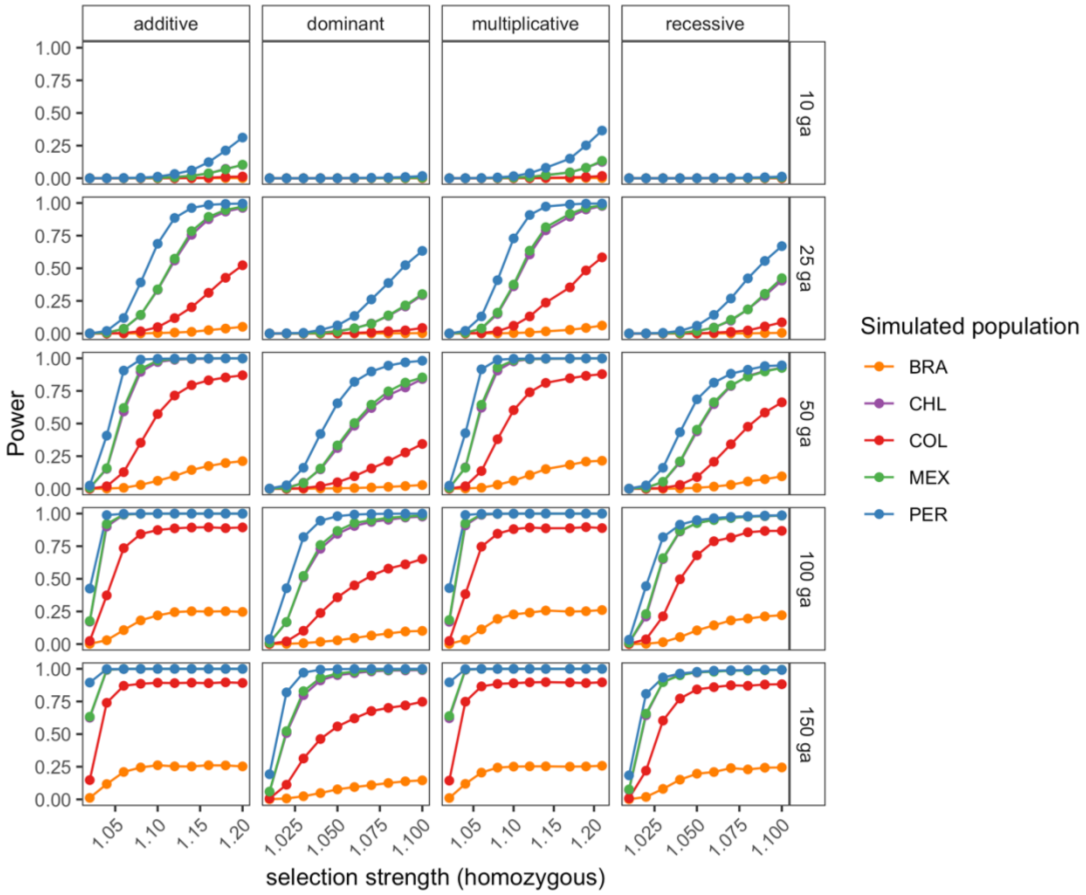


**Supplementary figure S2. Performance of AdaptMix to detect selection in the Native American source population of simulated Latin American populations.** Power to detect selection post-admixture in simulated admixed Latin American populations under three different admixture models, and four different selection models. See supplementary figure S1 legend for further details.

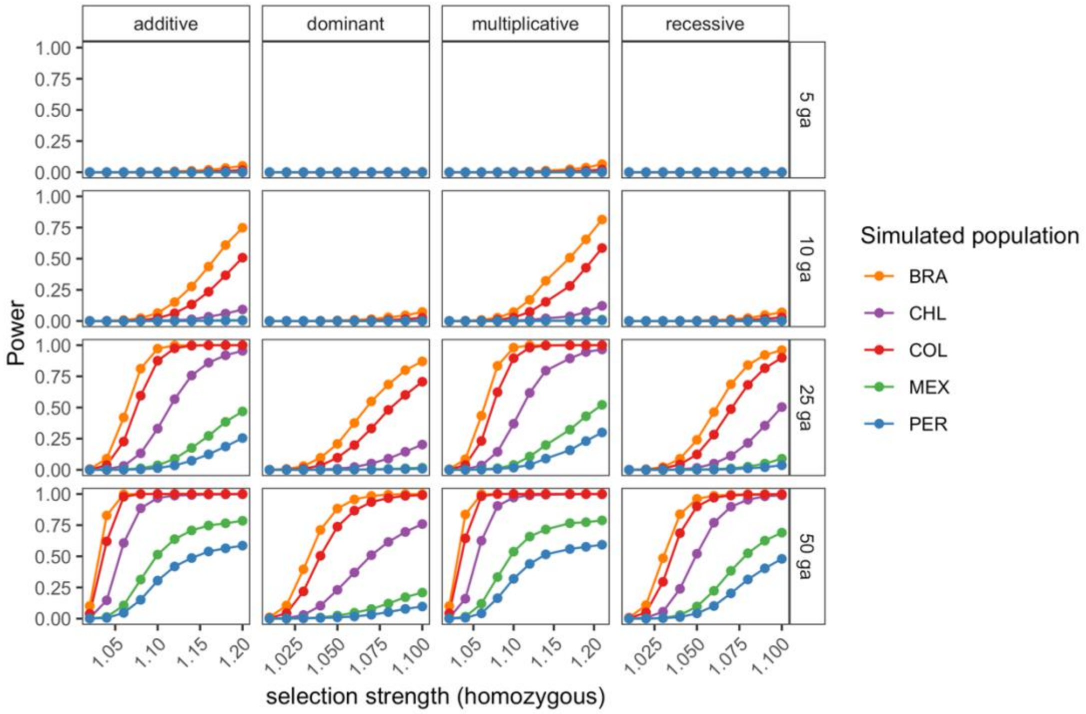


**Supplementary figure S3. Performance of AdaptMix to detect selection in the European source population of simulated Latin American populations.** Power to detect selection post-admixture in simulated admixed Latin American populations under three different admixture models, and four different selection models. See supplementary figure S1 legend for further details.

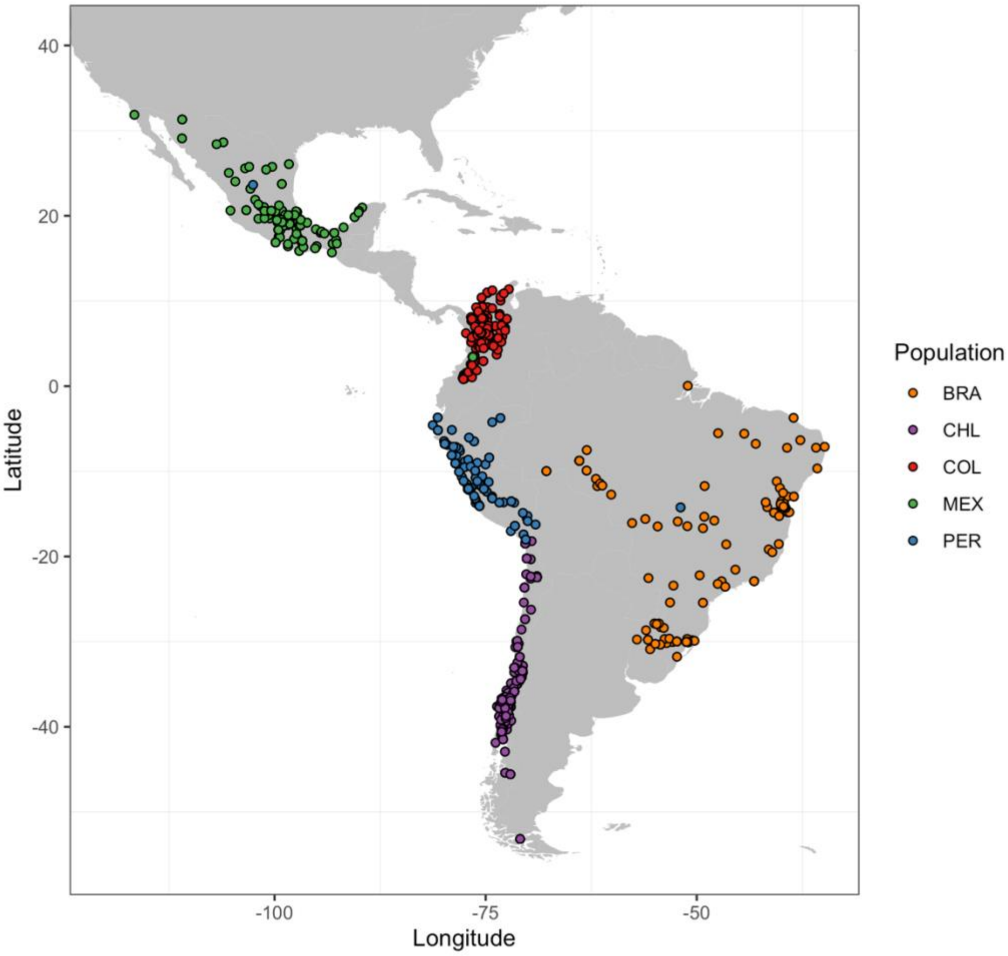


**Supplementary figure S4. Map of sampled CANDELA individuals.** Circles correspond to unique birthplaces.

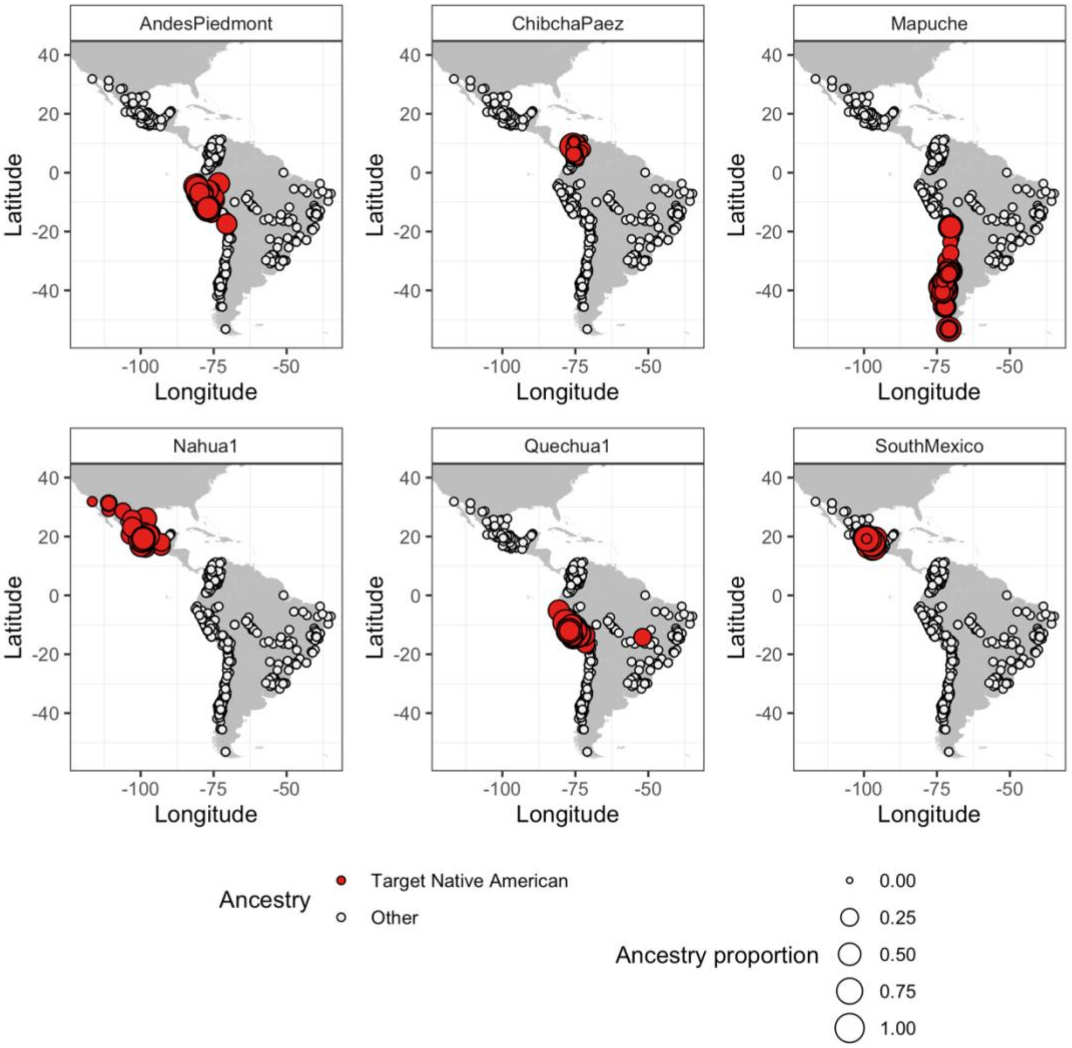


**Supplementary figure S5. Map of sampled CANDELA individuals based on their inferred ancestry matching to Native American reference groups.** Circle locations correspond to unique birthplaces, with red highlighting individuals with inferred Native American ancestry predominantly from the given source. The size of each red circle corresponds to the proportion of inferred ancestry matching to this source.

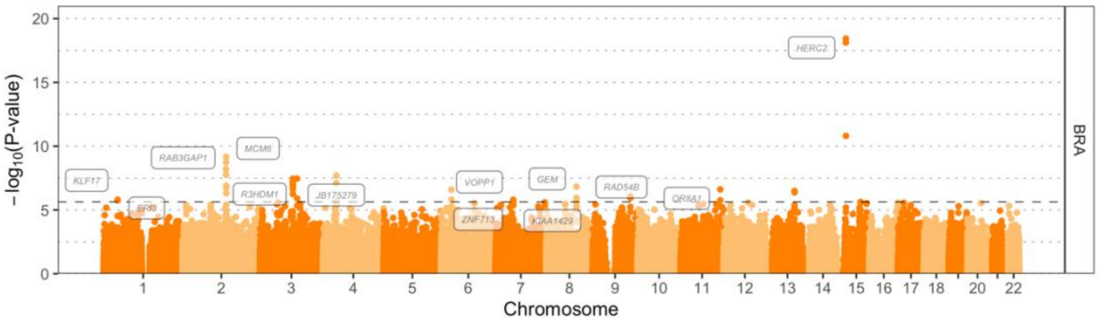


**Supplementary figure S6. Manhattan plot in the Brazilian population using GBR from the 1000 Genomes Project as donor European population.** The dashed horizontal lines indicate corresponds to the 99.99$^{th}$ percentile. Names of genes associated with SNPs above the 99.99$^{th}$ percentile cutoff are shown.

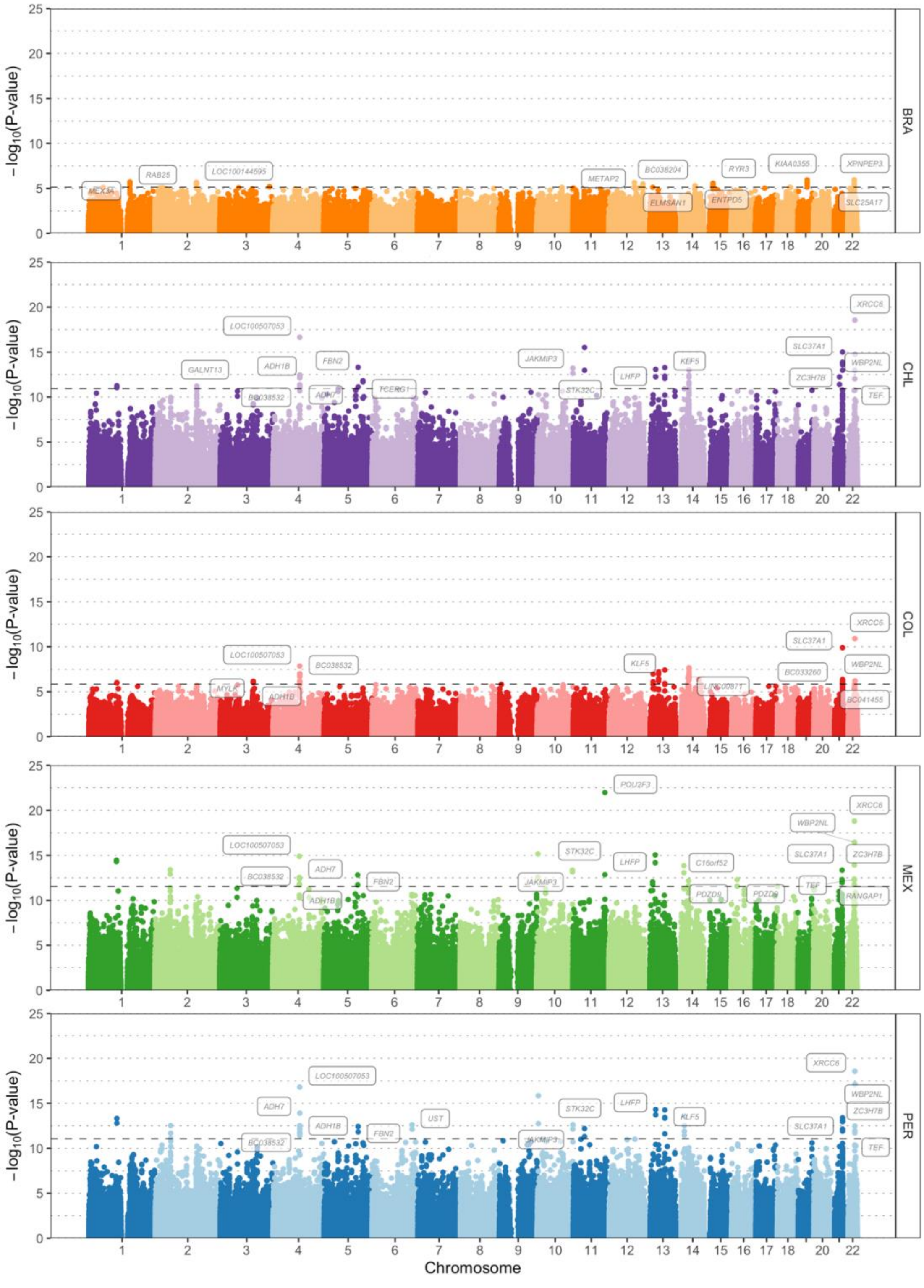


**Supplementary figure S7. Manhattan plots in the CANDELA cohort using CHB from the 1000 Genomes Project as donor Native American population.** The dashed horizontal lines indicate corresponds to the 99.99$^{th}$ percentile. Names of genes associated with SNPs above the 99.99$^{th}$ percentile cutoff are shown.

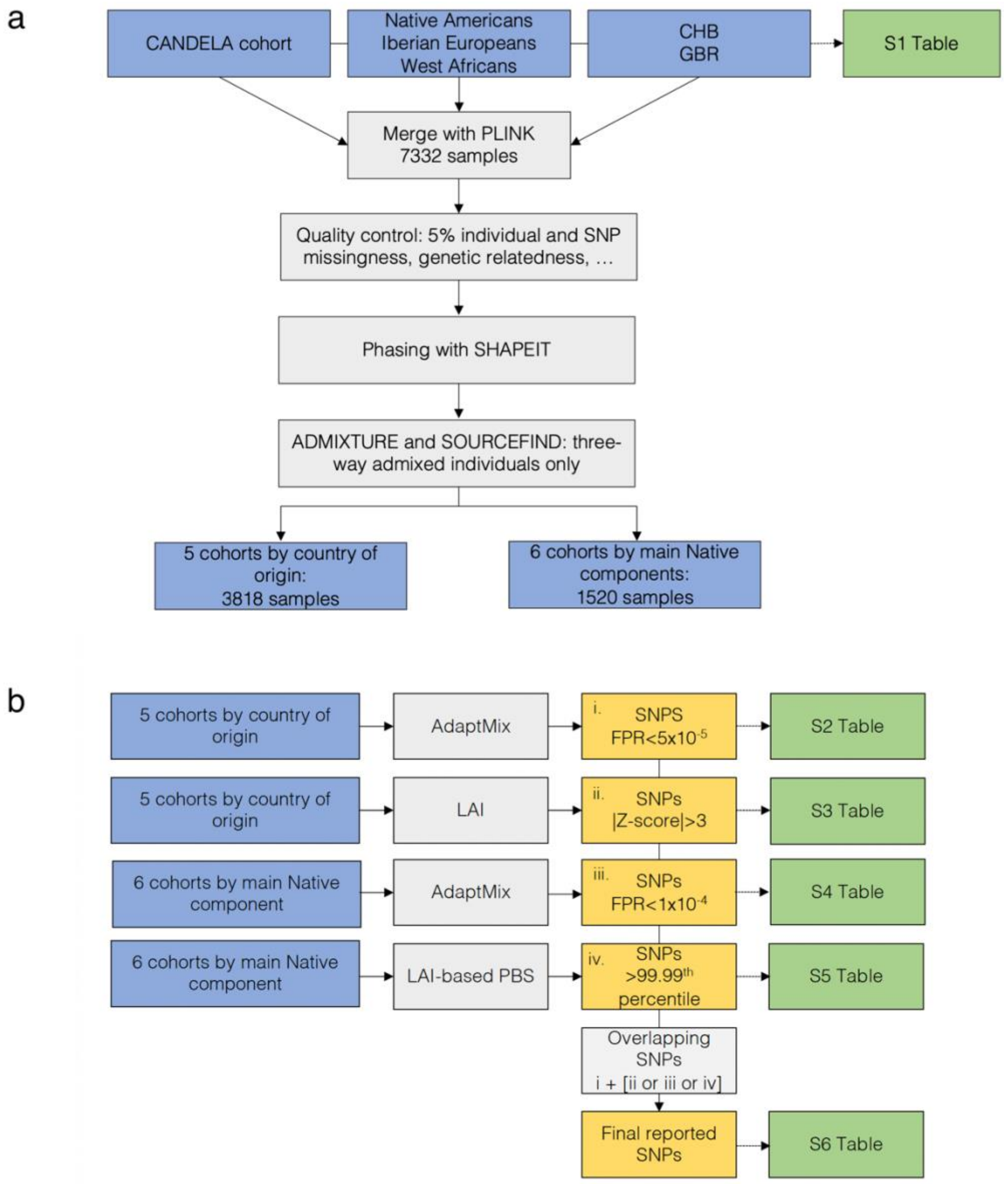


**Supplementary figure S8. Flowchart of analyses.**

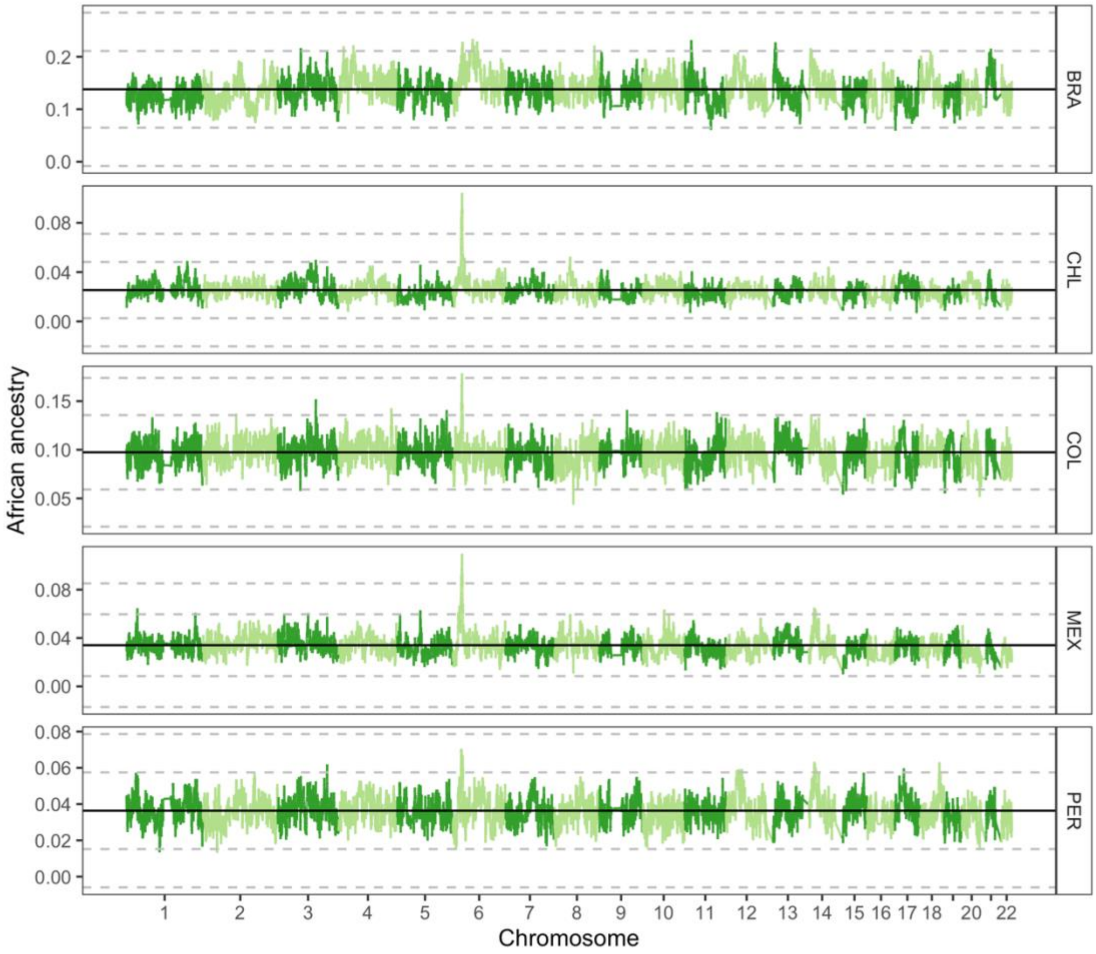


**Supplementary figure S9. African local ancestry in Latin American populations.** The proportion of African ancestry at each genomic location is shown. Solid black line shows the genome-wide average. Dashed grey lines show 3, and 6 standard deviations

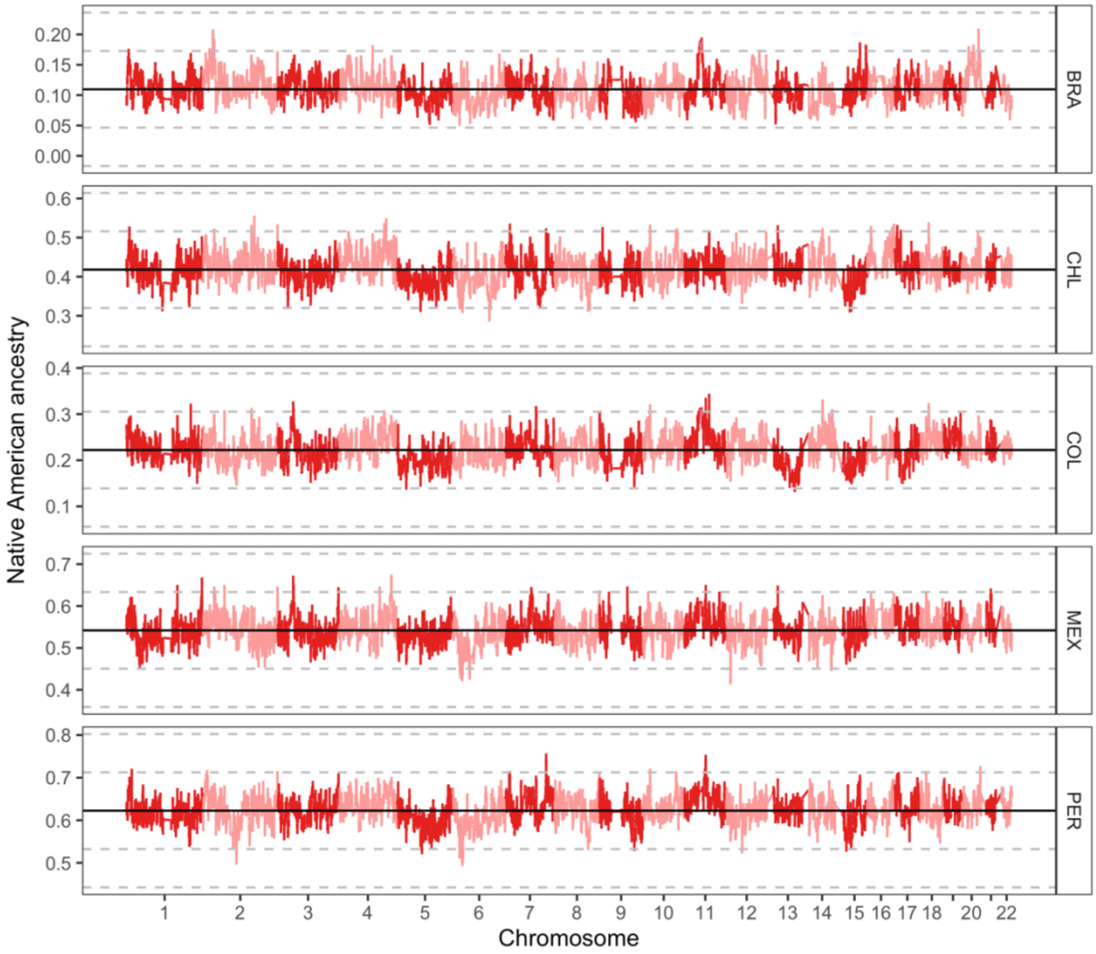


**Supplementary figure S10. Native American local ancestry in Latin American populations.** The proportion of Native American ancestry at each genomic location is shown. Solid black line shows the genome-wide average. Dashed grey lines show 3, and 6 standard deviations

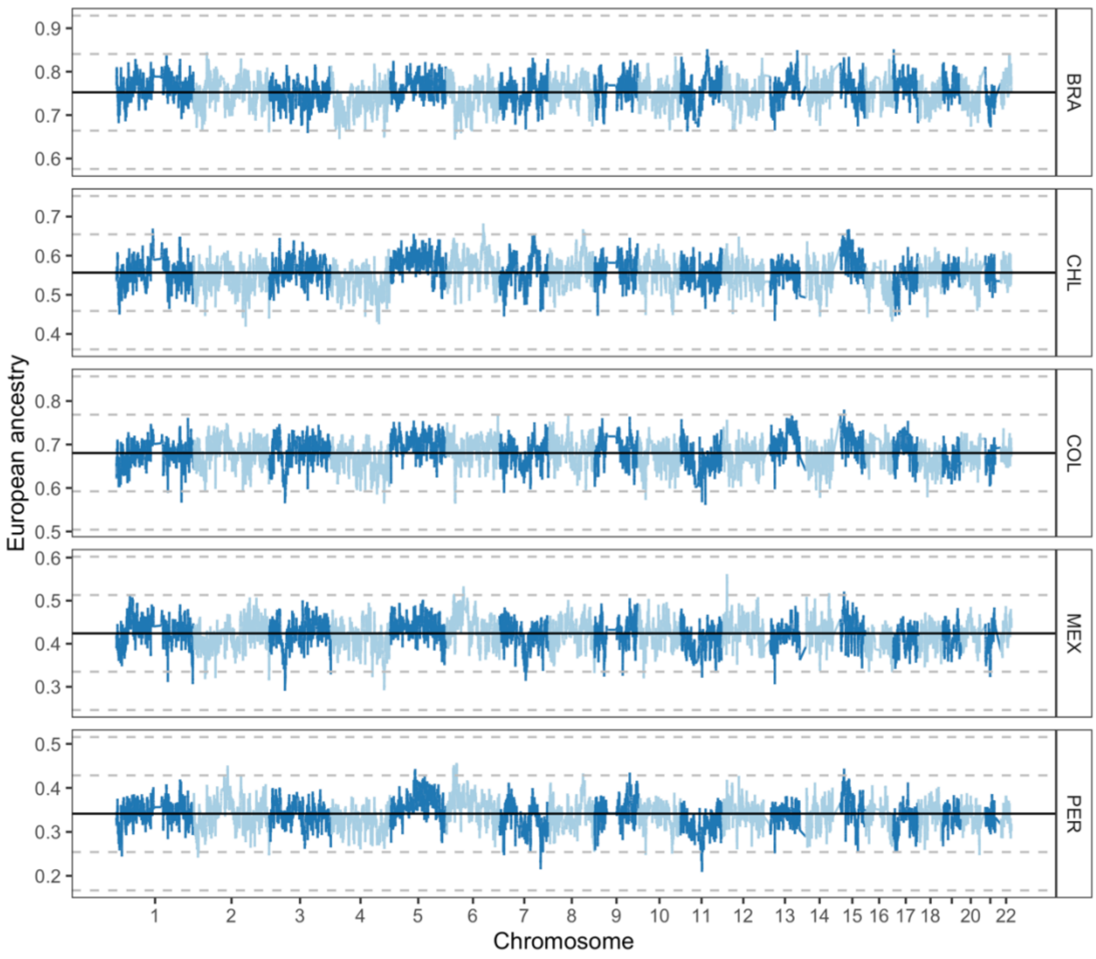


**Supplementary figure S11. European local ancestry in Latin American populations.** The proportion of European ancestry at each genomic location is shown. Solid black line shows the genome-wide average. Dashed grey lines show 3, and 6 standard deviations

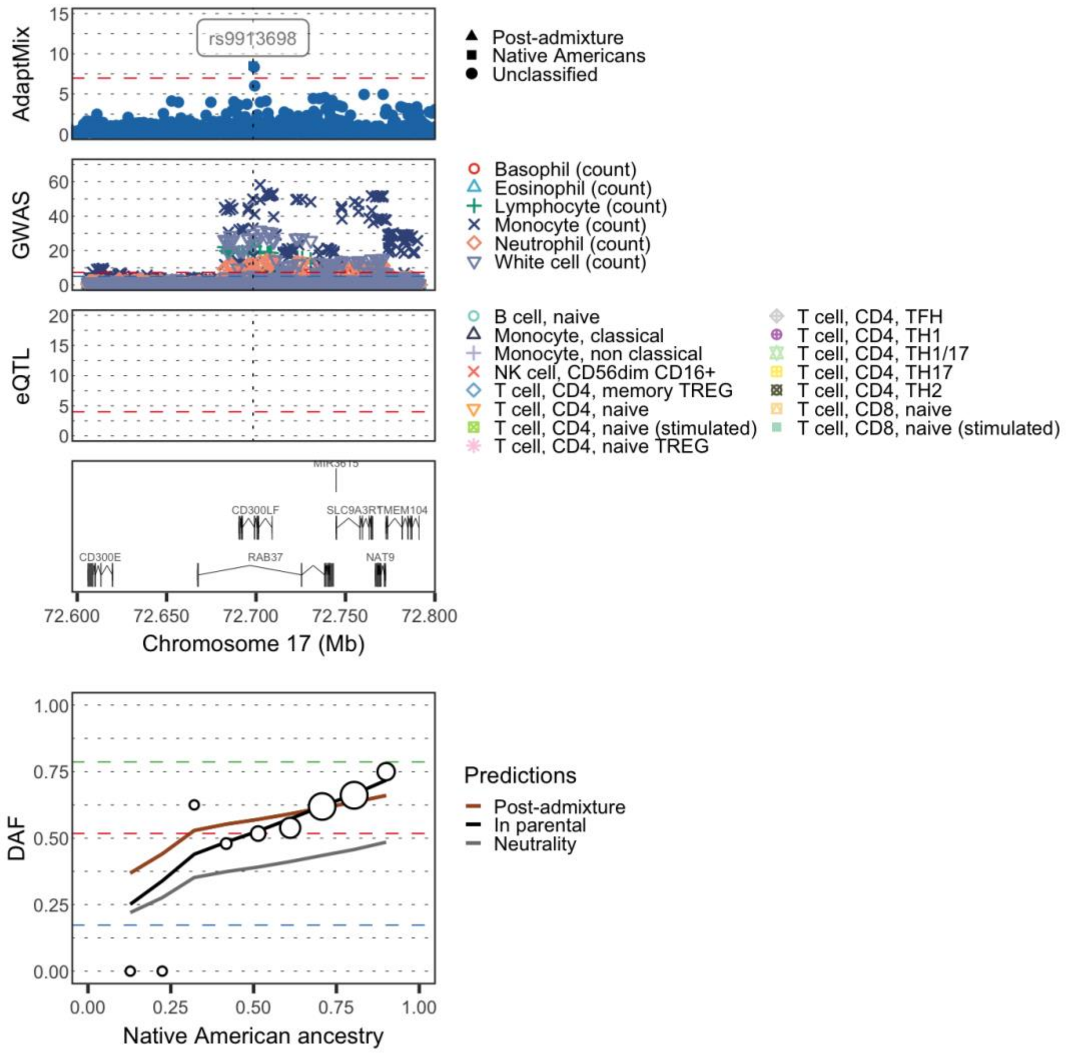


**Supplementary figure S12. Regional selection plot at the *CD300LF* gene in the Peruvian population.** The first four rows consist of $-\log_{10}$($P$-values) of SNPs: (row 1) from AdaptMix; (row 2) from a GWAS for immune-related cell counts[1]; (row 3) associated with gene expression of *CD300LF* from the DICE eQTL study[2]; with (row 4) depicting genes in the region (in Mb, build hg19 as reference). Horizontal dashed lines give significance thresholds of (row 1) $P$-value = $1\times10^{-5}$ (row 2) $P$-value = $1\times10^{-5}$ (blue line) and $P$-value = $5\times10^{-8}$ (red line), and (row 3) $P$-value = $1\times10^{-4}$. The lower plot depicts derived allele frequencies (DAF) in the Peruvian cohort (white circles) stratified by proportion of inferred Native American ancestry, for the SNP rs9913698. The sizes of the circles are proportional to the number of individuals in that particular bin. Lines give expected DAF under neutrality (grey), post-admixture selection (brown) or selection in the Native source (black). Horizontal dashed red, blue, and green lines depict DAF for surrogates to Native American, European, and African sources, respectively.

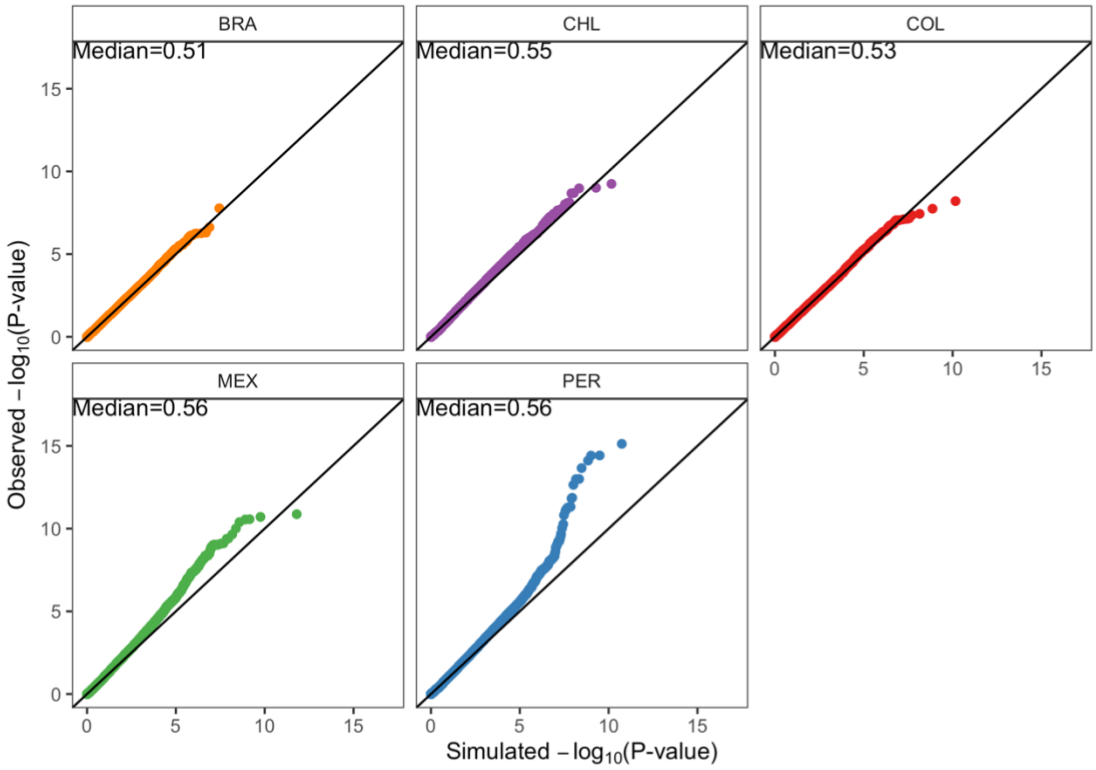


**Supplementary figure S13. Quantile-Quantile plot for AdaptMix in the CANDELA cohort**, comparing -$\log_{10}$(*P*-values) for observed SNPs versus -$\log_{10}$(*P*-values) for matching SNPs simulated under neutrality. Observed median *P*-values are provided on the upper left corner of each facet.

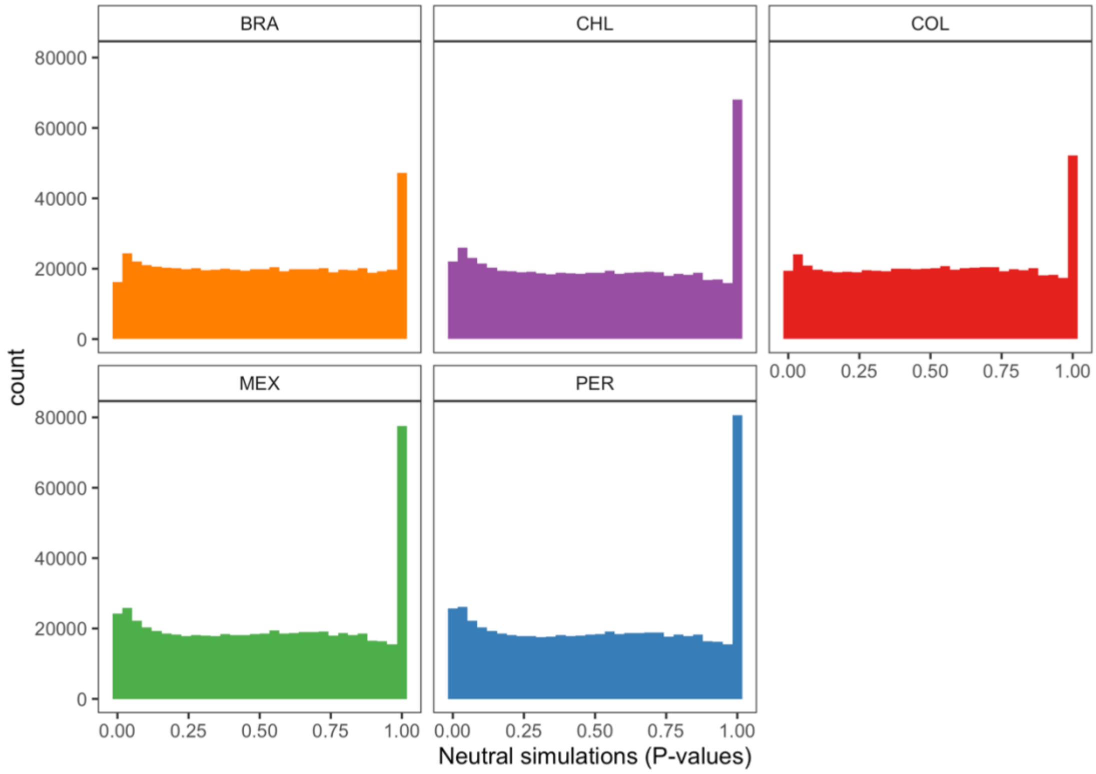


**Supplementary figure S14. Distribution of *P*-values under neutrality in simulated Latin American populations.**

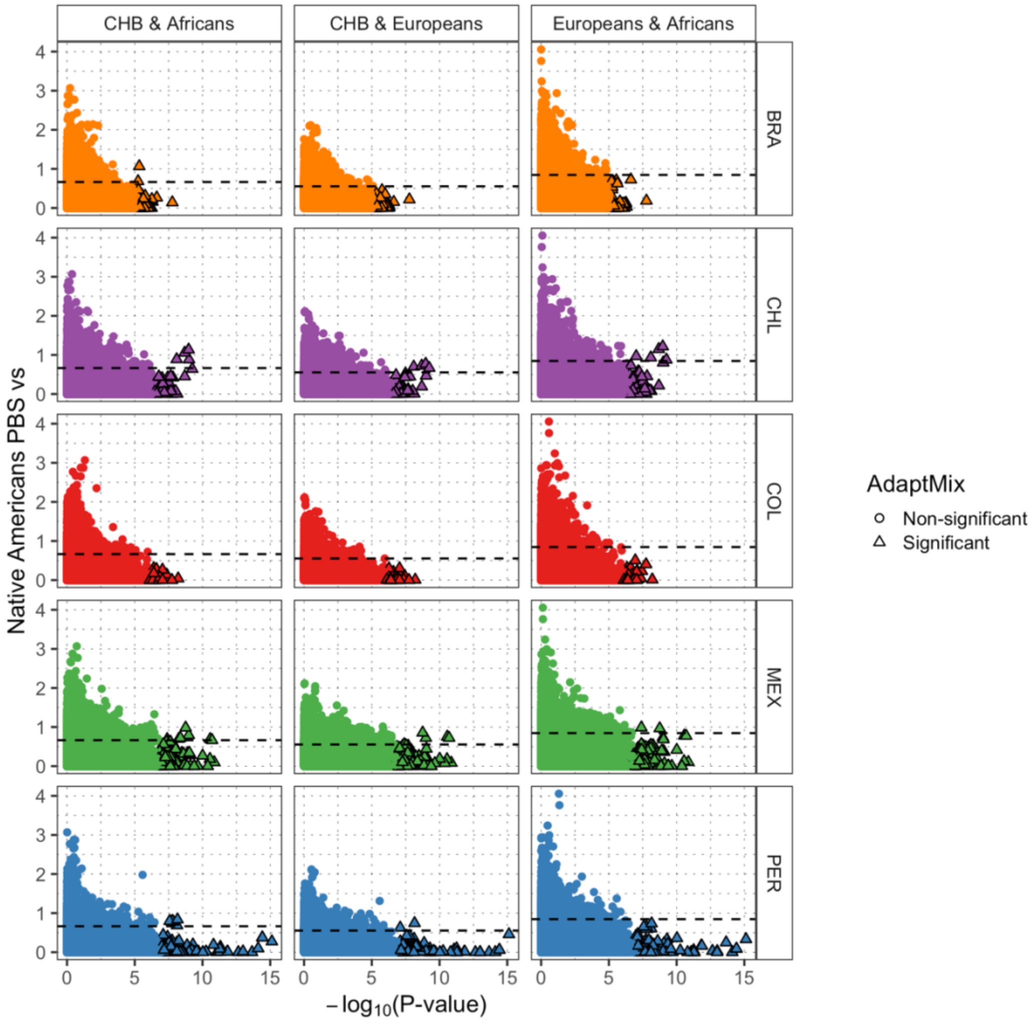


**Supplementary figure S15. Correlation between PBS (y-axis) and AdaptMix (x-axis) scores in the CANDELA cohort.** Black dashed line shows the $99^{th}$ percentile of PBS scores.

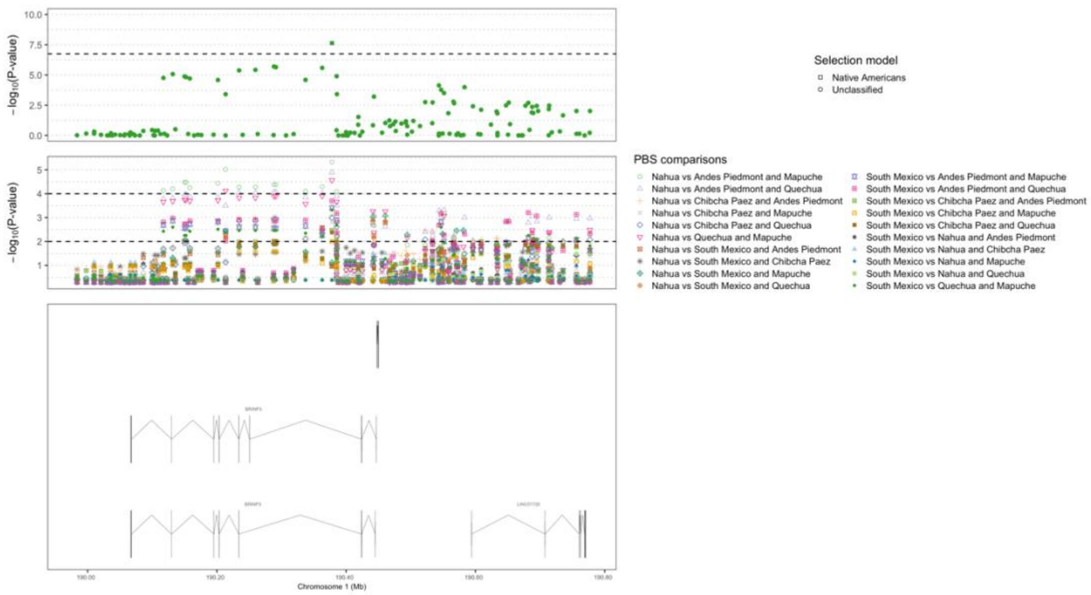


**Supplementary figure S16. Regional AdaptMix selection plot at 1q31 in the Mexican population.** The upper plot shows the $-\log_{10}$(*P*-values) of SNPs from AdaptMix, the middle plot shows the $-\log_{10}$(empirical *P*-values) of SNPs from PBS analysis, and the bottom plots shows the genes (including transcripts) in the region (in Mb, build hg19 as reference). The dashed lines refer to *P*-value cutoff that resulted in a 0.05% false-positive rate in neutral simulations for AdaptMix and the 99$^{th}$ and 99.99$^{th}$ percentiles for PBS.

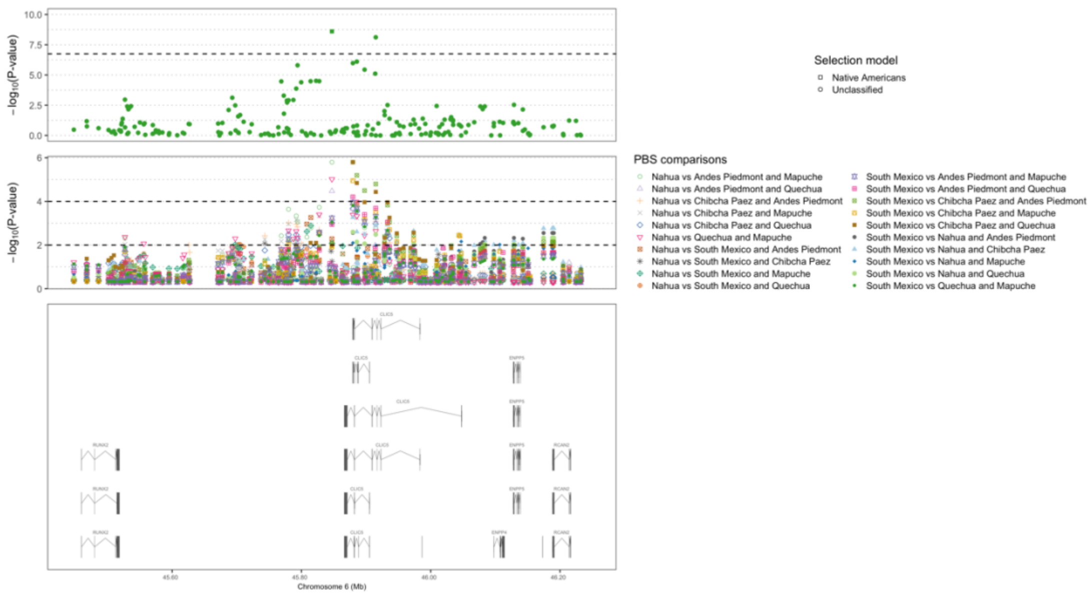


**Supplementary figure S17. Regional AdaptMix selection plot at 6p21 in the Mexican population.** See legend for supplementary figure S16 for details.

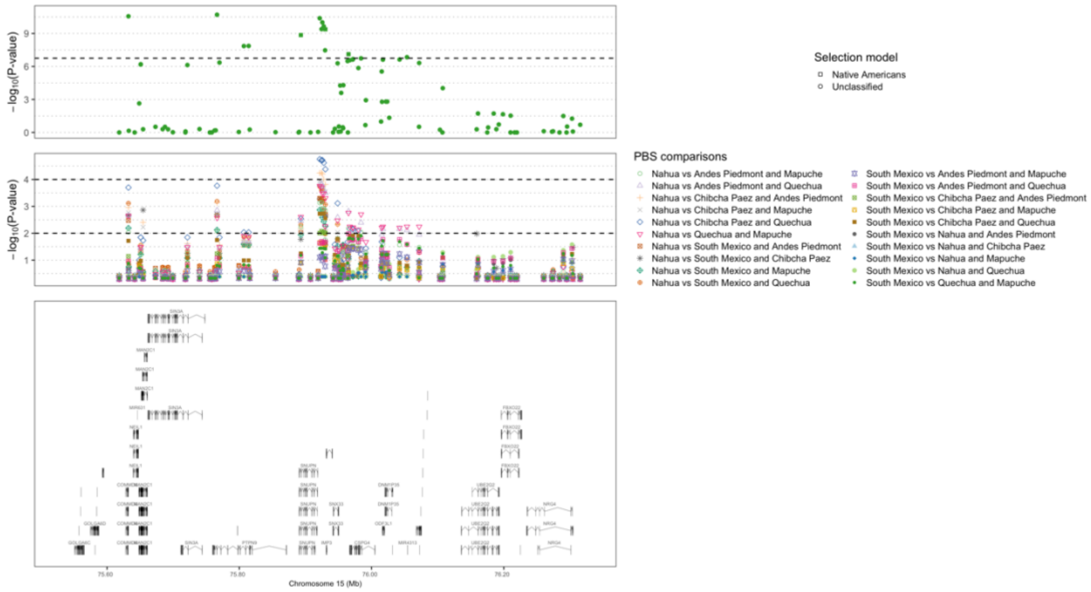


**Supplementary figure S18. Regional AdaptMix selection plot at 15q24 in the Mexican population.** See legend for supplementary figure S16 for details.

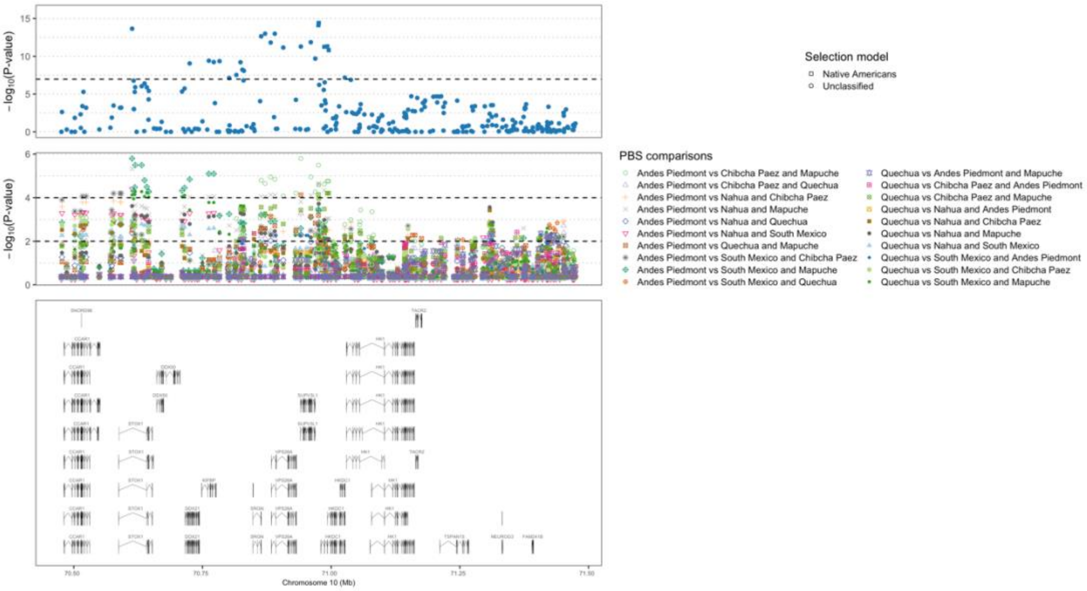


**Supplementary figure S19. Regional AdaptMix selection plot at 10q22 in the Peruvian population.** See legend for supplementary figure S16 for details.

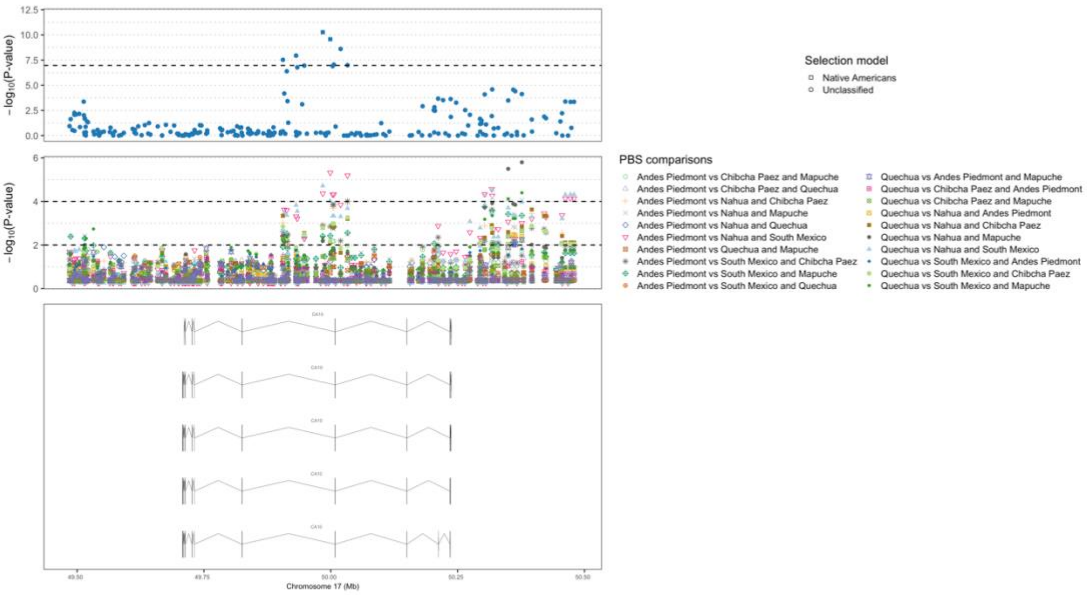


**Supplementary figure S20. Regional AdaptMix selection plot at 17q21 in the Peruvian population.** See legend for supplementary figure S16 for details.

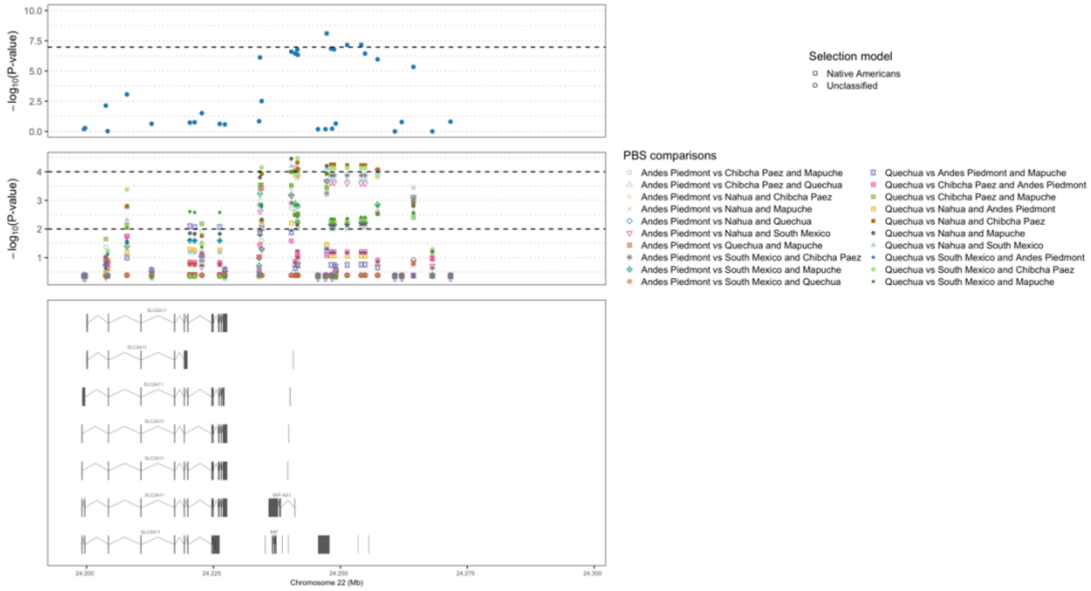


**Supplementary figure S21. Regional AdaptMix selection plot at 22q11 in the Peruvian population.** See legend for supplementary figure S16 for details.

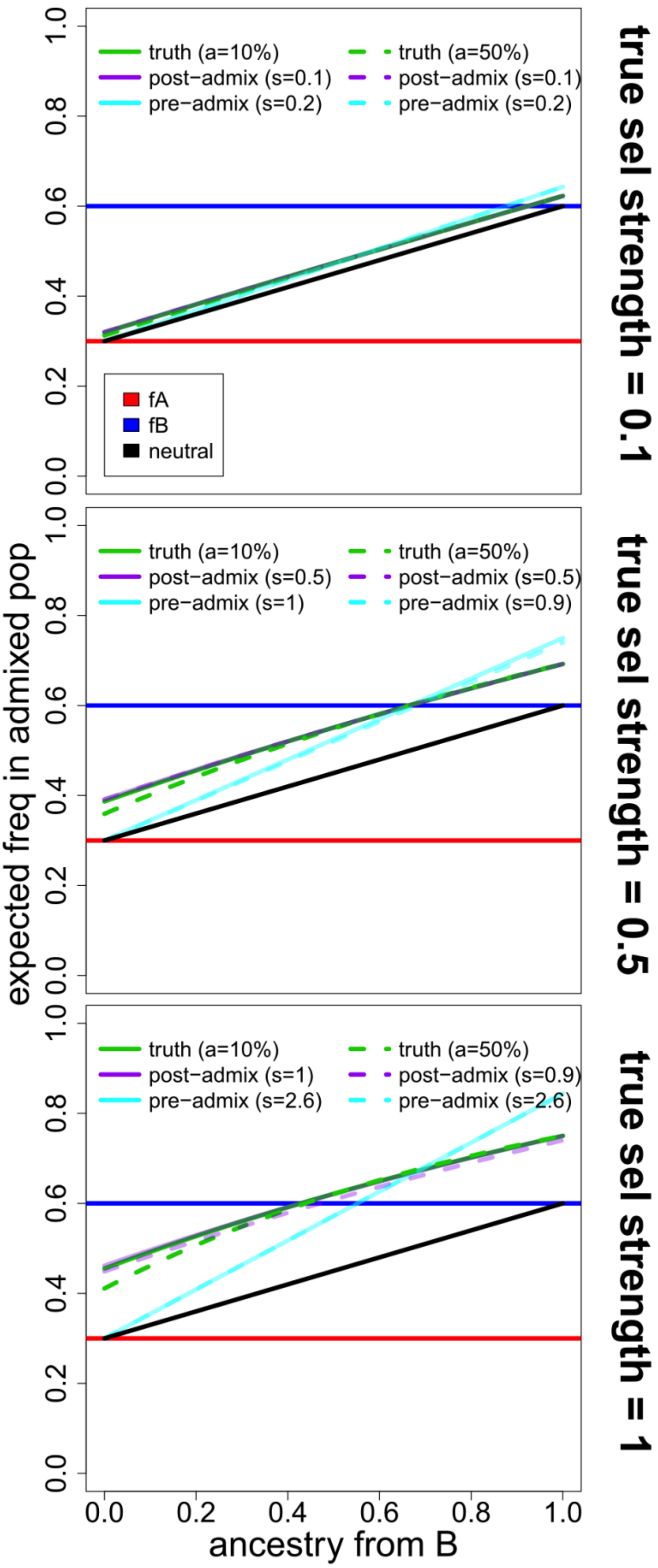

true sel strength = 0.1
truth (a=10%)
post−admix (s=0.1)
pre−admix (s=0.2)
truth (a=50%)
post−admix (s=0.1)
pre−admix (s=0.2)
fA
fB
neutral
true sel strength = 0.5
truth (a=10%)
post−admix (s=0.5)
pre−admix (s=1)
truth (a=50%)
post−admix (s=0.5)
pre−admix (s=0.9)
true sel strength = 1
truth (a=10%)
post−admix (s=1)
pre−admix (s=2.6)
truth (a=50%)
post−admix (s=0.9)
pre−admix (s=2.6)
expected freq in admixed pop
ancestry from B
0.0
0.2
0.4
0.6
0.8
1.0

**Supplementary figure S22. Expected allele frequencies (green) at a SNP versus admixture received from source B, in a population undergoing selection following admixture between sources A and B.** The total selection strengths summed across generations (under a multiplicative model) are shown on the right-hand side of each panel. However, a={10%,50%} of the migrants from A are not subjected to this selection, mimicking new migrants from A. The allele frequencies of population A and B are given by the fA and fB horizontal lines, respectively. The fit of the neutral model is given with the black line. Also depicted is the theoretical best fit of selection occurring post-admixture (purple lines) or in source B prior to admixture (cyan lines), assuming a multiplicative selection model. The best-fitting selection coefficient ($s$) under each of the two models is given in parentheses in the legend, representing the value of $s$ in $[0, 0.1, \ldots, 2.9, 3]$ that had lowest mean-squared-error with the truth. Note that for total selection strengths ≥0.5, visually the post-admixture selection better fits than the data than pre-admixture selection, despite the influx of new migrants from A that have not been subjected to this selection pressure.

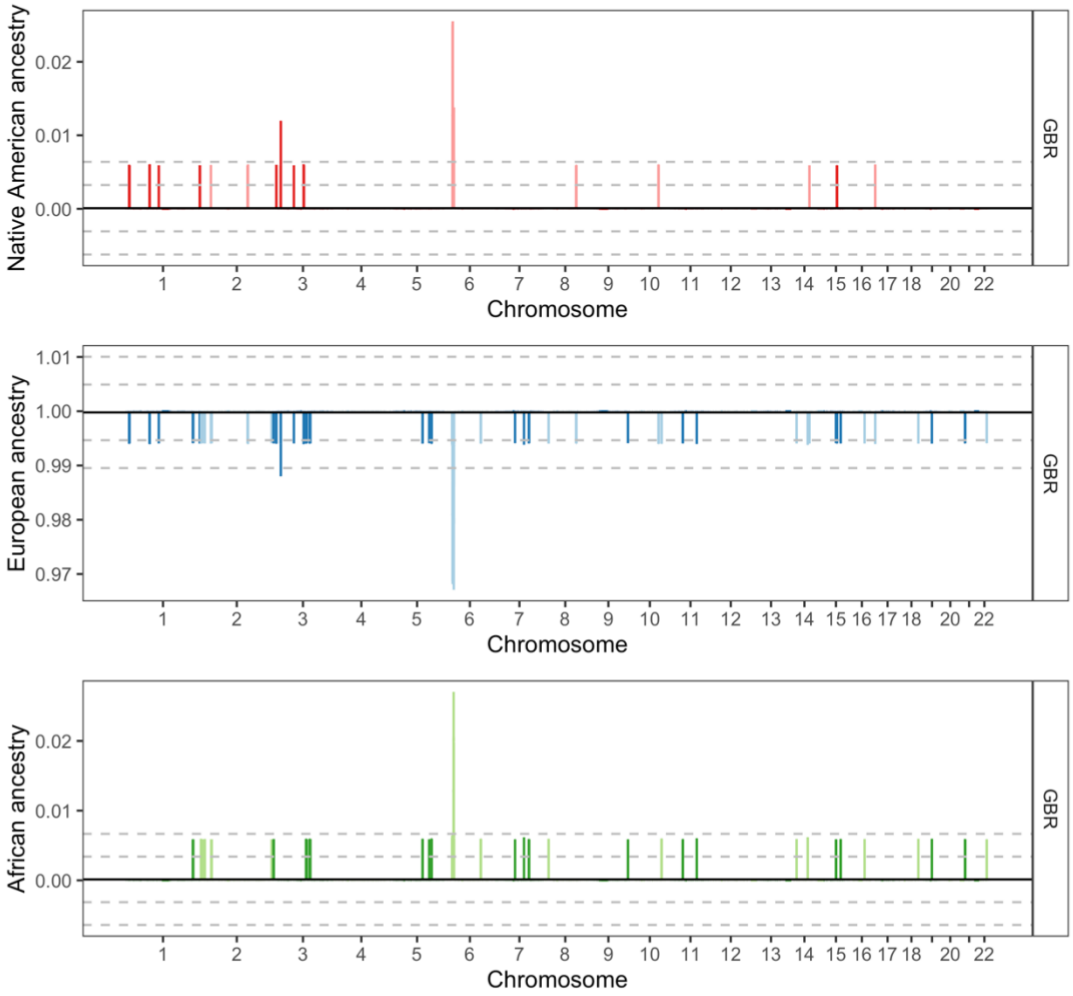


**Supplementary figure S23. Local ancestry deviations in the GBR population from the 1000 Genomes Project.** The proportion of Native American (in red), European (in blue), and African ancestry (in greed) at each genomic location is shown. Solid black line shows the genome-wide average. Dashed grey lines show 3, and 6 standard deviations.

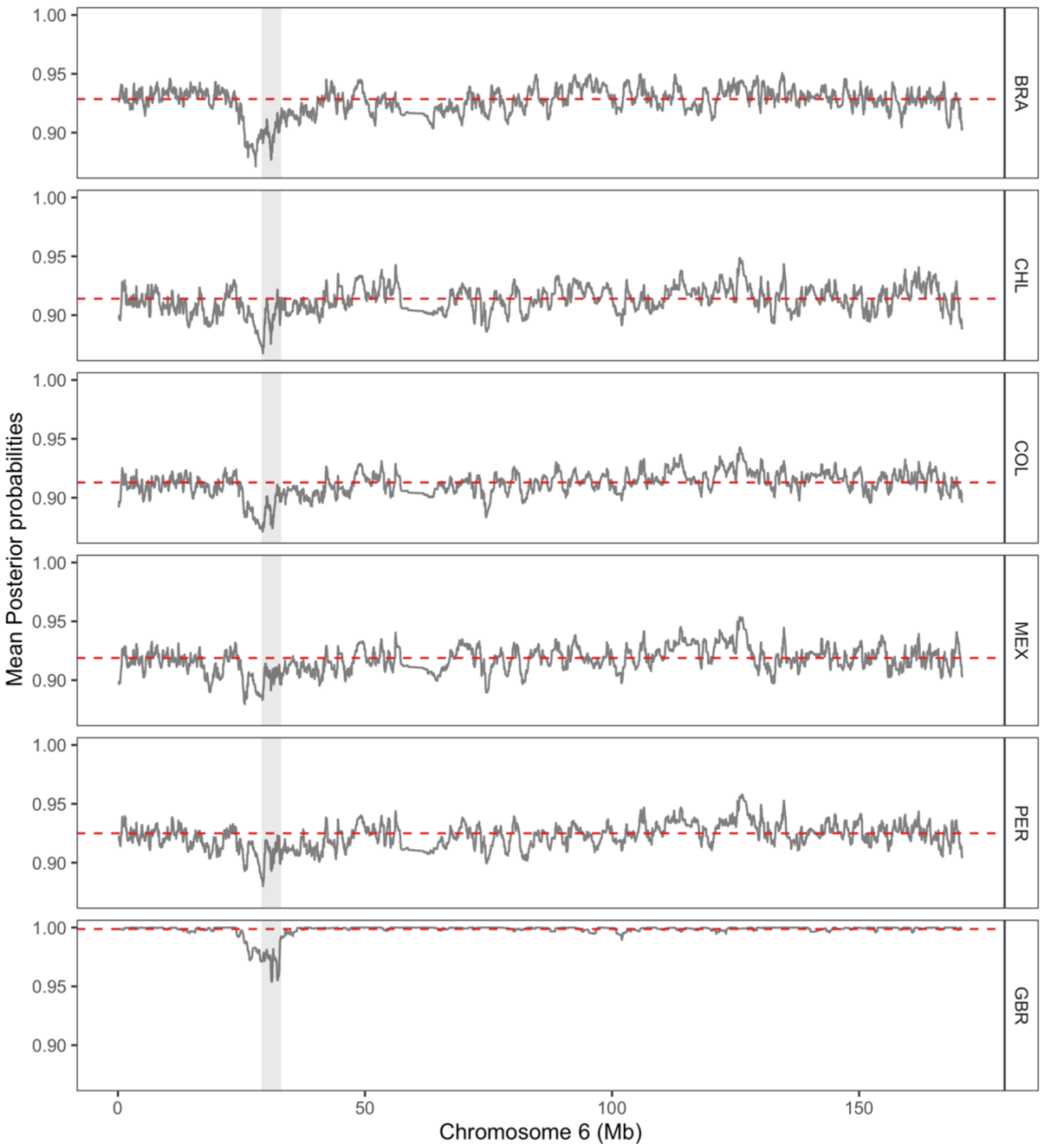


**Supplementary figure S24. Mean posterior probabilities of local ancestry assignments in the CANDELA cohort and GBR population from the 1000 Genomes Project.** Mean posterior probabilities for chromosome 6 are shown as grey lines. The region of lowest mean posterior probabilities corresponds to the HLA region highlighted in grey. The mean genome-wide posterior probabilities are shown as red dashed lines.

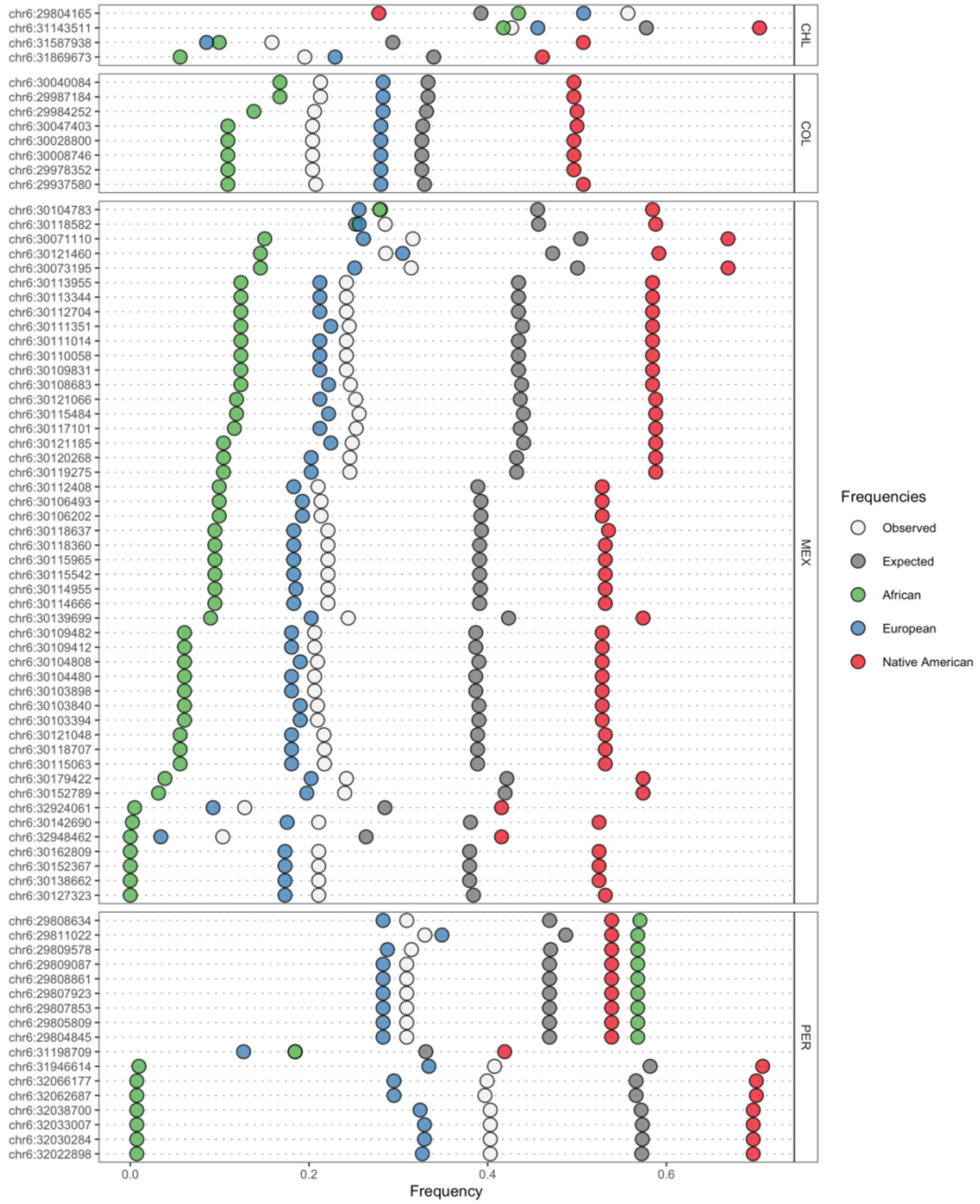


**Supplementary figure S25. Allele frequencies at significant SNPs at HLA loci.** Rows are ordered by decreasing allele frequency in the African surrogate group. Expected is the allele frequency based on neutrality under our model, i.e. assuming each CANDELA cohort's frequency can be described as a linear mixture of the surrogates' frequencies.

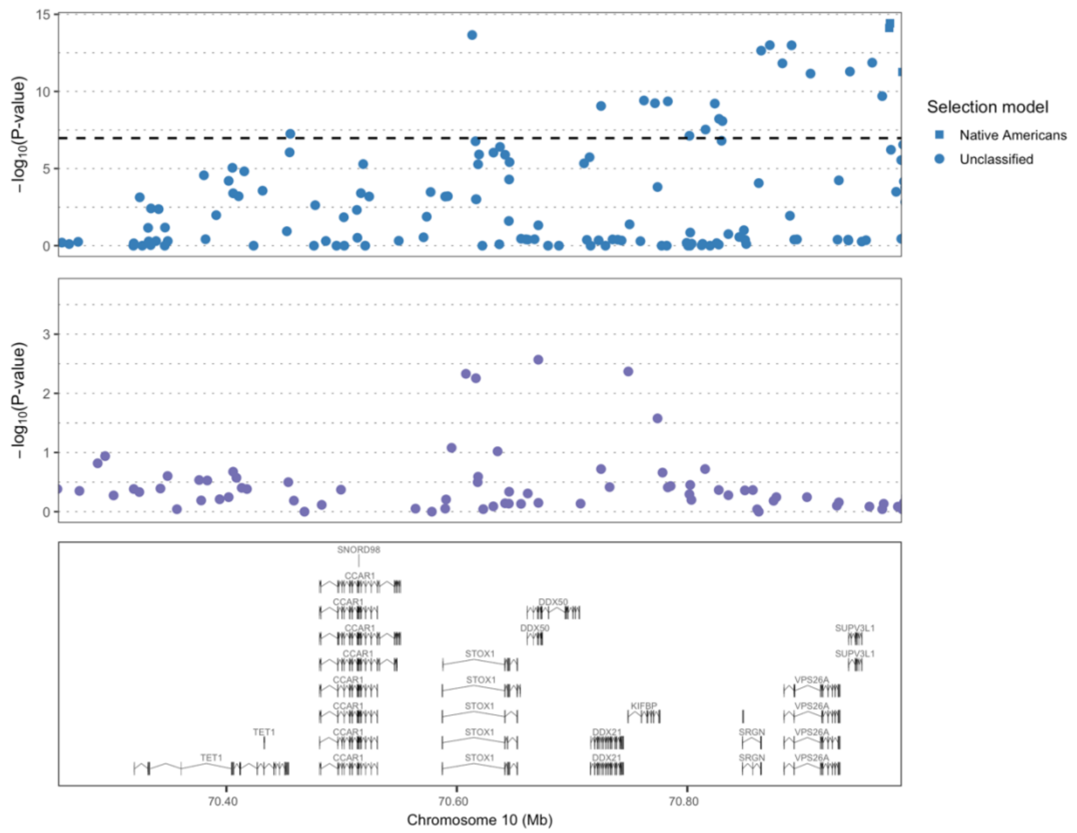


**Supplementary figure S26. Selection signals in the Peruvian cohort and transmission disequilibrium test for preeclampsia at *STOX1*.** The upper plot shows the $-\log_{10}$(*P*-values) of SNPs from AdaptMix in the Peruvian cohort. The dashed lines refer to *P*-value cutoff that resulted in a $5\times10^{-5}$ false-positive rate in neutral simulations for AdaptMix. The middle plot shows the $-\log_{10}$(*P*-values) of SNPs associated to preeclampsia in an Andean cohort based on the Badillo Rivera and Nieves-Colón et al (2021) study. The bottom plot shows the genes (including transcripts) in the region (in Mb, build hg19 as reference).

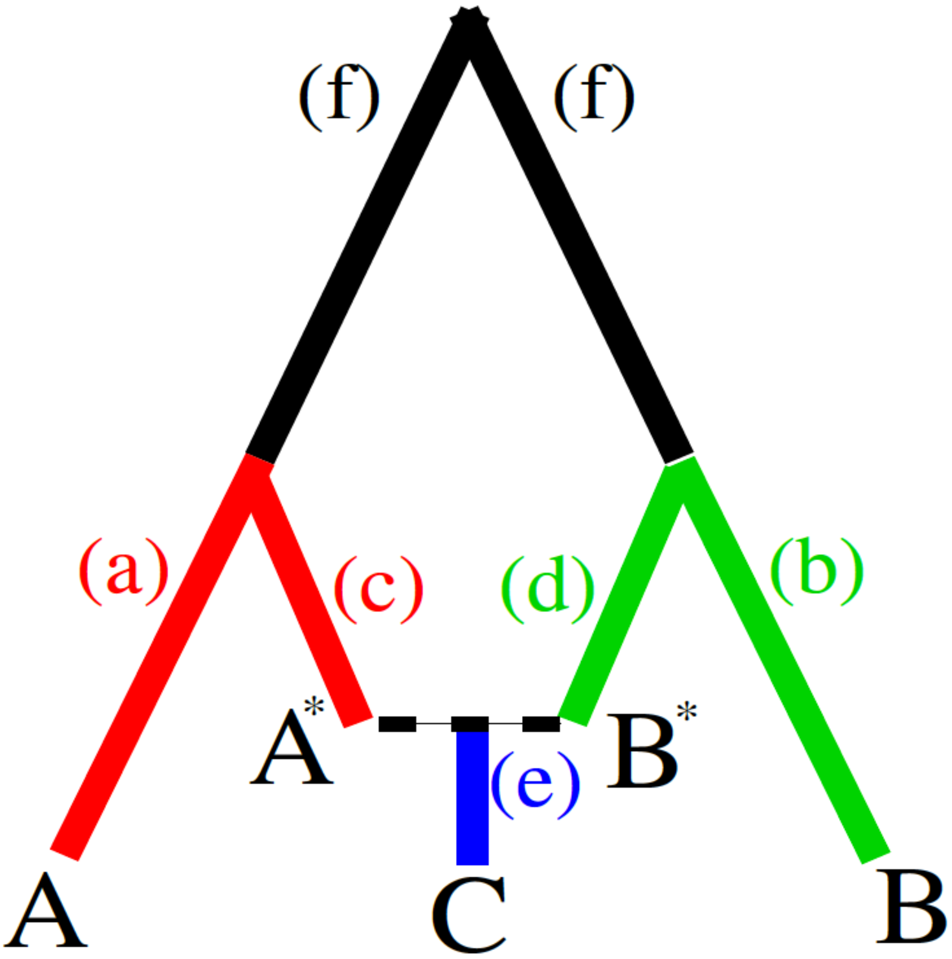


**Supplementary figure S27. An admixture graph showing the relationship between two parental populations, and an admixed population.** Tree relating sampled populations A, B, and C, with C descending from admixture between unsampled populations $A^*$ and $B^*$.
